# Textural features for pathway-level representation of omics data in biological networks

**DOI:** 10.64898/2026.07.12.737672

**Authors:** Andrey Alexeyenko

## Abstract

More than 50 years ago, Haralick and co-authors proposed a family of gray-level co-occurrence statistics that became known as textural features. These features are widely used in image analysis, but their application to biological networks has remained limited because cellular networks are sparse, irregular graphs rather than regular pixel grids.

This work presents a network-adapted version of Haralick texture analysis for generating pathway-level features from gene-level omics profiles. The resulting profiles reduce dimensionality and can be used as candidate predictors of anti-cancer drug response. Performance of these features is compared with original gene expression variables and with pathway features from network enrichment analysis (NEA), whose robustness has been demonstrated previously. Although technically simpler than NEA, Haralick features showed comparable sensitivity. More importantly, selected Haralick features were preserved between *in vitro* drug screens and clinical treatment-associated survival analyses, supporting their potential use for prioritizing robust pathway-level drug-response correlates.

## Introduction

Pathway enrichment analysis, used to summarize individual molecular profiles at the pathway level, is a popular approach in omics data interpretation. It helps characterize sets of genes that are differentially expressed or otherwise altered, for example in pathological conditions (Huang et al. 2009). It can be particularly useful for characterizing individual samples when pathway scores provide an alternative to original gene-level omics profiles, an approach initially presented as single-sample gene set enrichment analysis, ssGSEA (Barbie et al. 2009). This can reduce dimensionality from thousands of genes to hundreds, or fewer, pathways. In settings where pathological alteration events are individually sparse or poorly detectable, summarizing them across samples and patients can be beneficial, especially in cancer, which is known for extreme heterogeneity at all molecular levels (Heng 2015).

Pathway features can be evaluated in individual samples by accounting for the functional state of their member genes. However, a more common and simpler approach has been to analyze gene set enrichment, GSEA (or overrepresentation, ORA), i.e. quantifying overlap between genes in a pathway and genes that distinguish a biological condition (Subramanian et al. 2005) (Huang et al.2009).

Modern human cellular networks, for example those derived from FunCoup (Alexeyenko and Sonnhammer 2009), STRING (Szklarczyk et al. 2015), or PathwayCommons (Rodchenkov et al.2020), contain much more information than databases of discrete pathways. Such networks can include millions of edges, mostly supported by experimental evidence but not confined within pathway limits, and can therefore provide a richer biological context for sample characterization. We previously proposed network enrichment analysis (NEA) (Alexeyenko et al. 2012) (Jeggari and Alexeyenko 2017), which evaluates the overrepresentation of network links between pathway genes and sample-specific genes, or altered gene sets (AGSs) rather than shared genes as in ORA. NEA is more complex than ORA because it uses a global interaction network and accounts for an additional parameter, the node degree of each gene. This brought higher sensitivity and greater feature robustness. It is worth noting that global networks probably have high false positive and false negative rates: many declared links do not exist, while many existing links remain unknown. However, this problem is equally relevant to network-free methods, which may include irrelevant genes in pathways or false positives from omics datasets. In both cases, the purpose of enrichment analysis is to evaluate signal strength against a noisy background. Enrichment analysis is facilitated by the availability of multiple expert-curated pathway databases and ontologies. Many functional gene sets (FGSs) provided by such resources could, either by design or by coincidence, be relevant to biological functions, conditions, and diseases. This makes the identified correlates not only statistically significant, but also biologically interpretable.

A more mechanistic alternative to enrichment would be to characterize pathway functional states, perturbations, or damage by monitoring the activity of key genes. However, pathway-state quantification, as in SPIA (Tarca et al. 2009), PARADIGM (Vaske et al. 2010), Pathifier (Drier et al. 2013), PROGENy (Schubert et al. 2018), or VIPER (Alvarez et al. 2016), is limited either by the requirement to know pathway maps and interaction modes precisely or by ignoring the broader and deeper network context available from multi-source resources such as those mentioned above.

The present work implements and analyzes a trade-off between pathway enrichment and pathway state approaches, while accounting for the global cellular network context.

Texture features were proposed by Haralick and co-authors in 1973 (Haralick et al. 1973) and have been widely used to analyze pixel maps of satellite images (Moya et al. 2019), wood patterns(Kobayashi et al. 2015), biological tissues, and cellular compartments (Brynolfsson et al. 2017). To my knowledge, Barker-Clarke and co-authors were the first, and so far the only team to apply Haralick textural features to biological network analysis. Using a collection of simpler example graphs, they showed that the features expectedly varied with graph topology (Barker-Clarke et al.2022).

Here, Haralick texture features are tested for their ability to capture clinically relevant molecular landscapes. The statistical criterion is whether pathway-level correlates of clinical phenotypes can be detected and then preserved across experimental and clinical datasets. Molecular marker discovery is a major use case for omics data, but cancer correlates are often poorly reproducible because relevant molecular events are heterogeneous and sparse at the individual-gene level.

Moving from genes, mutations, methylation events, or expression values to pathways can reduce dimensionality by summarizing sparse events into interpretable functional units. NEA applications have shown that network context can improve sensitivity and robustness compared with network-free enrichment methods (Alexeyenko et al. 2012) (Franco et al. 2019) (Brink et al. 2019) (Alexeyenko et al. 2022). As with PCA, pathway-level profiling reduces the number of variables and may limit overfitting in downstream multivariate models; unlike PCA, it preserves biological interpretability and facilitates visualization.

The usability of *in vitro* models of drug response has been repeatedly questioned because molecular landscapes of cancer cell lines were found to deviate substantially from those of primary tumors (Domcke et al. 2013). Although a more comprehensive analysis demonstrated overall consistency of molecular aberrations between primary tumors and relevant cell lines (Iorio et al. 2016), the clinical relevance of the discovered *in vitro* correlates was not investigated. A discouraging comparison between two *in vitro* screens was also published (Haibe-Kains et al.2013), although later work showed that it was not statistically impeccable (Franco et al. 2019). The latter work also demonstrated that concerns about cancer cell lines arise when individual gene profiles are considered and can be partly addressed at the pathway level. The performance of NEA-based features was comprehensively tested against existing pathway methods, both network-based and network-free (Franco et al. 2019). NEA exceeded the alternatives in statistical power, for example because ORA and related methods suffered from sparse set overlaps, and in the robustness of identified drug-sensitivity correlates across independent datasets and between *in vitro* and clinical response profiles. A technical complication in NEA was the need to delineate individual, sample-specific AGSs, which required optimization and fine-tuning of AGS derivation parameters for each new analytical setting. In this work, I use a similar framework to test the performance of textural features against original gene-expression profiles and NEA features. Comparisons with other enrichment techniques have been performed previously and are therefore not repeated here.

Unlike the example analyses of Barker-Clarke and co-authors, which were performed on whole networks, the present study is driven by the practical need of identifying clinically relevant and usable correlates. The pipeline therefore generates pathway-level texture features from pathway-specific subgraphs of the global human network. Sample-specific feature values are computed from gene-expression values assigned to pathway nodes. In parallel, NEA features are generated for the same pathway collection, and original gene-expression features are used as predictors in the same statistical models. The key comparison criteria are sensitivity, specificity, and reproducibility of drug-response correlates within and across *in vitro* and clinical datasets.

The compared methods are evaluated in single-feature models, with the expectation that the most robust features could later be used in multivariate predictors. Because multivariate prediction would require independent validation, this study focuses on the reproducibility of single-feature correlates. First, feature classes are compared in cancer cell-line data while controlling the false discovery rate. Next, identified correlates are tested for reproducibility in alternative *in vitro* screens. Finally, the most challenging comparison asks whether *in vitro* correlates are preserved in clinical follow-up data from patients treated with the same drugs.

## Results

In the first step of the algorithm, network node values are discretized, usually into *G*=2^*n*^ levels (Fig. 1A). Next, a *G* × *G* matrix of level co-occurrences is calculated for neighbor nodes.

**Figure 1.**
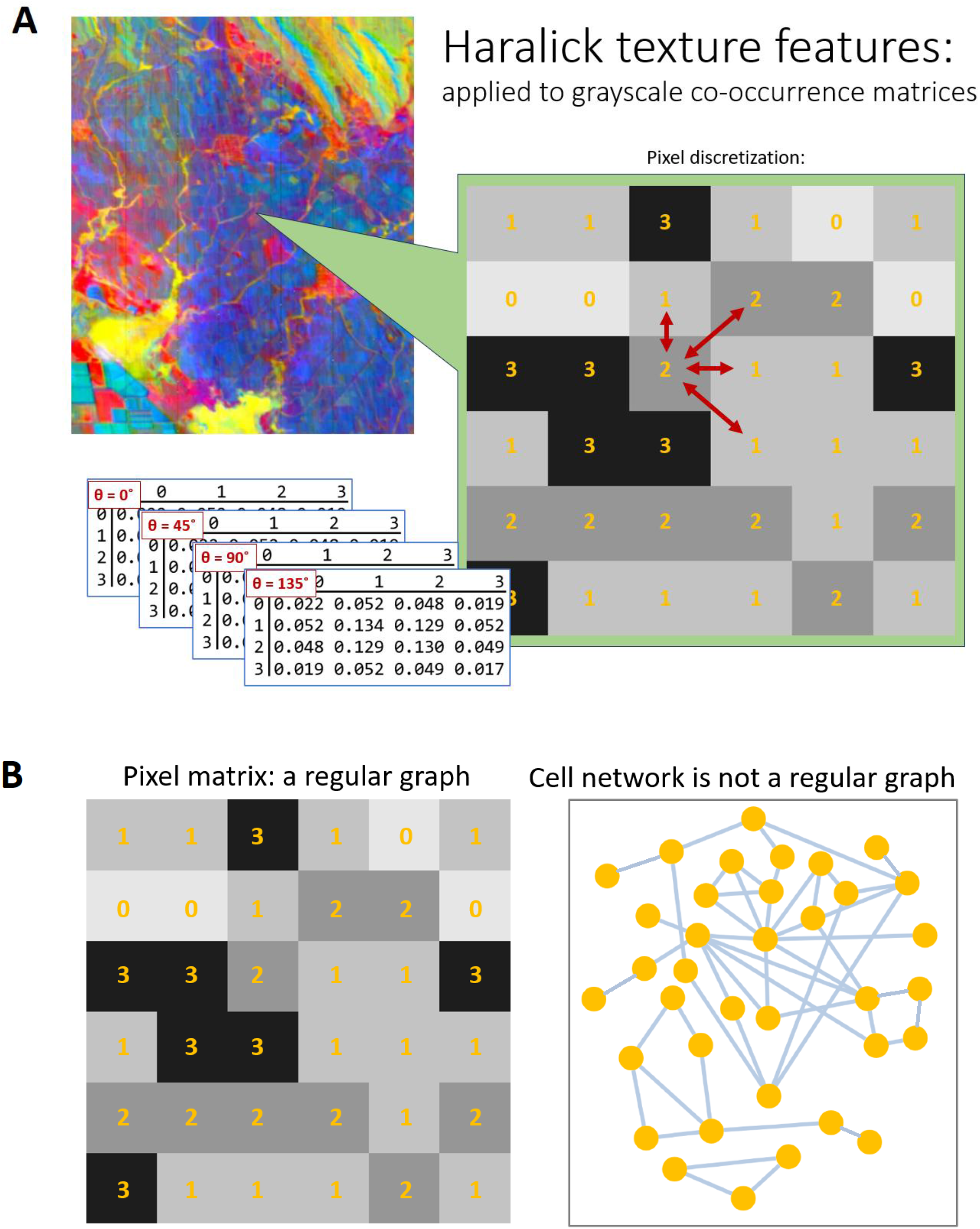
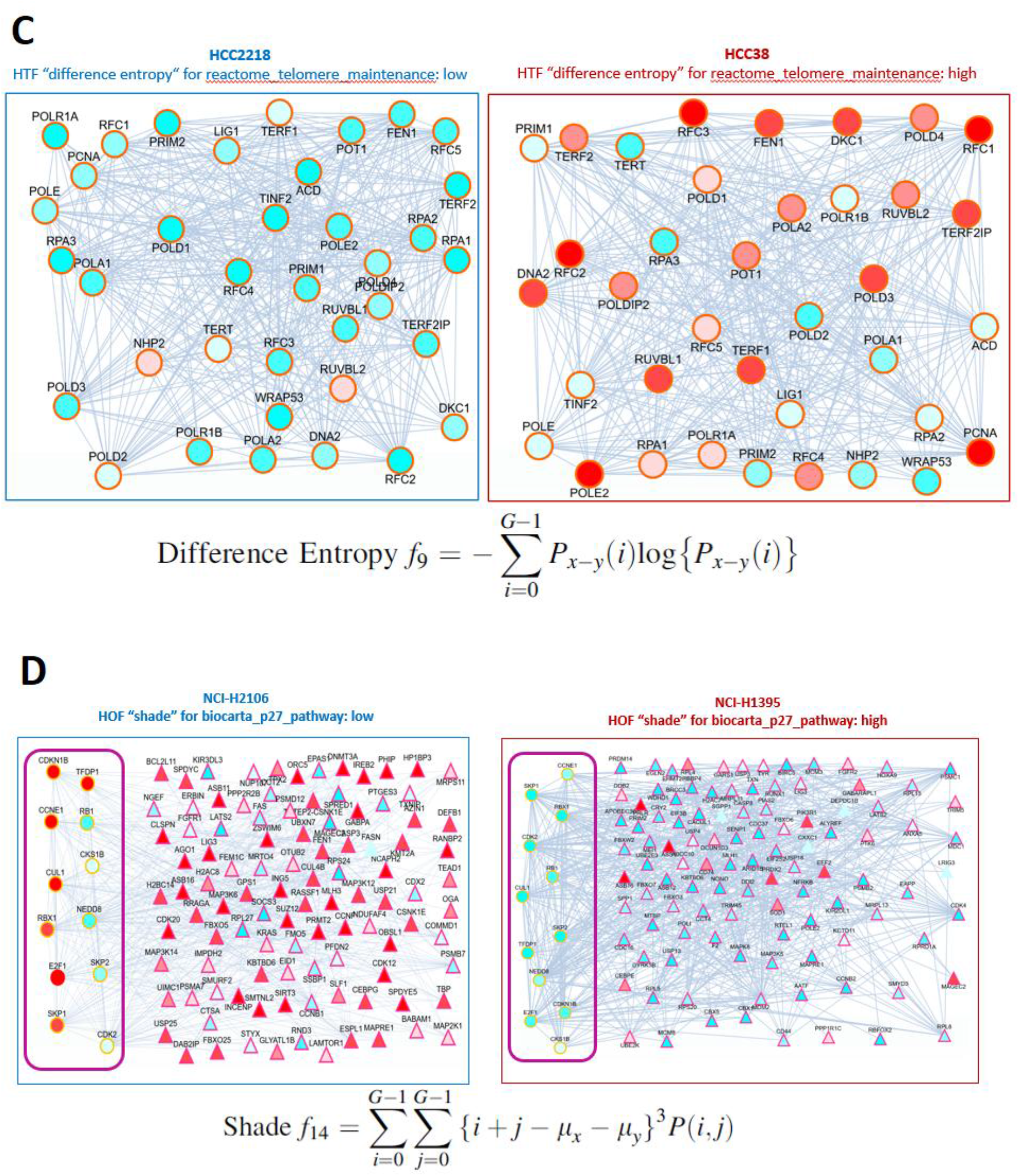
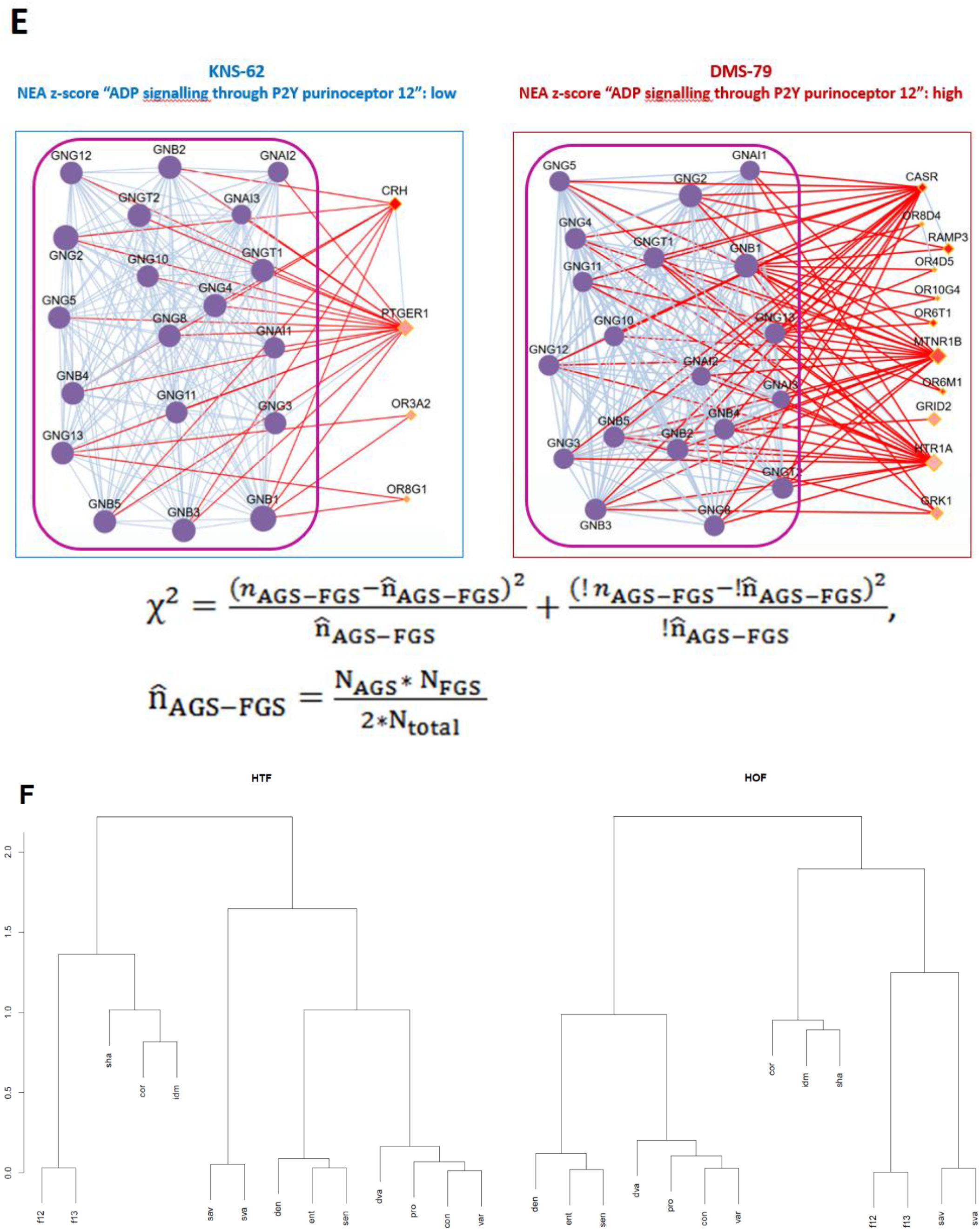
Outline of the Haralick texture calculation. A. A color image is converted to a strongly discretized (e.g. 2^2^ levels, 0…3) grayscale matrix. Probabilities of level adjacency are calculated for each of the four possible directions from 0°to 135°, if needed. B. Textural features are traditionally applied to regular graphs in which the number of neighbors per pixel is constant. Biological networks, by contrast, have unequal node degrees and are usually undirected, which makes accounting for directionality unnecessary. C. As an example, the HTF feature “difference entropy” (`den`) treats “activity-connectivity” patterns as the non-uniformity of gray-level differences between adjacent nodes, that is, genes connected by network links within a functional gene set (FGS). Within the FGS “Telomere maintenance pathway”, the cell line HCC38 received a higher `den` value because the activity levels of neighboring genes, shown as shades from blue to red on a continuous scale, tended to differ at certain ratios, mostly “low vs. medium” and “medium vs. high”. By contrast, in the cell line HCC2218 the levels were more uniform, resulting in a lower `den` value. D. In this example, the feature “shade” (`sha`) was calculated in HOF mode, i.e. accounting for both within- and outside-pathway genes (the former are surrounded by the magenta frames). For this feature, lower gray levels in adjacent nodes contributed to higher `sha` values. E. NEA estimates network connectivity between pathway genes and sample-specific sets of the most altered genes, in this example the top 50 genes most different from the collection mean in the given sample (diamonds in shades of pink). Since the activity of pathway genes is irrelevant here, the respective nodes (violet) are not shaded. Only links between AGS and FGS (red) are accounted for in the analysis. For statistical convenience, the resulting χ2 values are further converted to normally distributed z-scores. Gene node sizes reflect node degrees. F. Similarity of Haralick features characterizing pathway profiles of CCLE samples. Clustering was performed using Ward’s minimum variance method (“`ward.D2`” in the R function `hclust`) across feature profiles of samples in the CCLE expression dataset.

Topologically, raster images analyzed by Haralick’s method are regular graphs (Chen 1997), in which every image pixel has exactly four adjacent neighbors, or eight if diagonals are counted. Even if the analysis is expanded to more remote neighbors, graph regularity is preserved. A crucial difference in biological networks is that their node degrees are highly irregular (Fig. 1B), usually following power-law or lognormal distributions (Clauset et al. 2009). To accommodate this property, the original code from (Kobayashi et al. 2015;) was modified. Another difference is that neighboring pixels can optionally be matched at four alternative angles from 0°to 135°, whereas orientation is irrelevant for a network with undirected edges.

In this work, the analysis is restricted to pathway-specific subgraphs of the global network. In the default HTF mode, Haralick texture features are calculated only from edges connecting pathway-member genes. The alternative HOF mode additionally includes boundary edges in which only one of the two connected genes belongs to the analyzed pathway (Fig. 1C vs. 1D).

Unlike texture features, network enrichment analysis does not account directly for the activity of pathway genes (Fig. 1E). Instead, each cell line is characterized by an altered gene set (AGS). Network links between member genes in a pathway or functional gene set (FGS) of another type and genes in the AGS are then summed and normalized by values expected by chance, based on the cumulative node degrees, i.e. total numbers of links in the global network, of the AGS and FGS. Beyond trivial significance estimation, normalization gives higher importance to less connected, and hence potentially more context-specific, gene nodes. As an alternative to conventional multi-node FGSs, NEA can also consider single-node FGSs, treating individual genes as “potential pathways” (Franco et al. 2019). This option, GNEA, was also included in the comparative analysis.

### Initial comparative analysis and detection of drug-feature correlates *in vitro*

From the CCLE gene-expression dataset, the following feature categories were calculated for each cell-line sample (Table 1): log-normalized z-scores of original RNA-seq values (GE), Haralick texture features within pathways (HTF) and by including pathway boundaries (HOF), and network enrichment features at the pathway level (NEA) and individual gene-node level (GNEA).

**Table 1.**
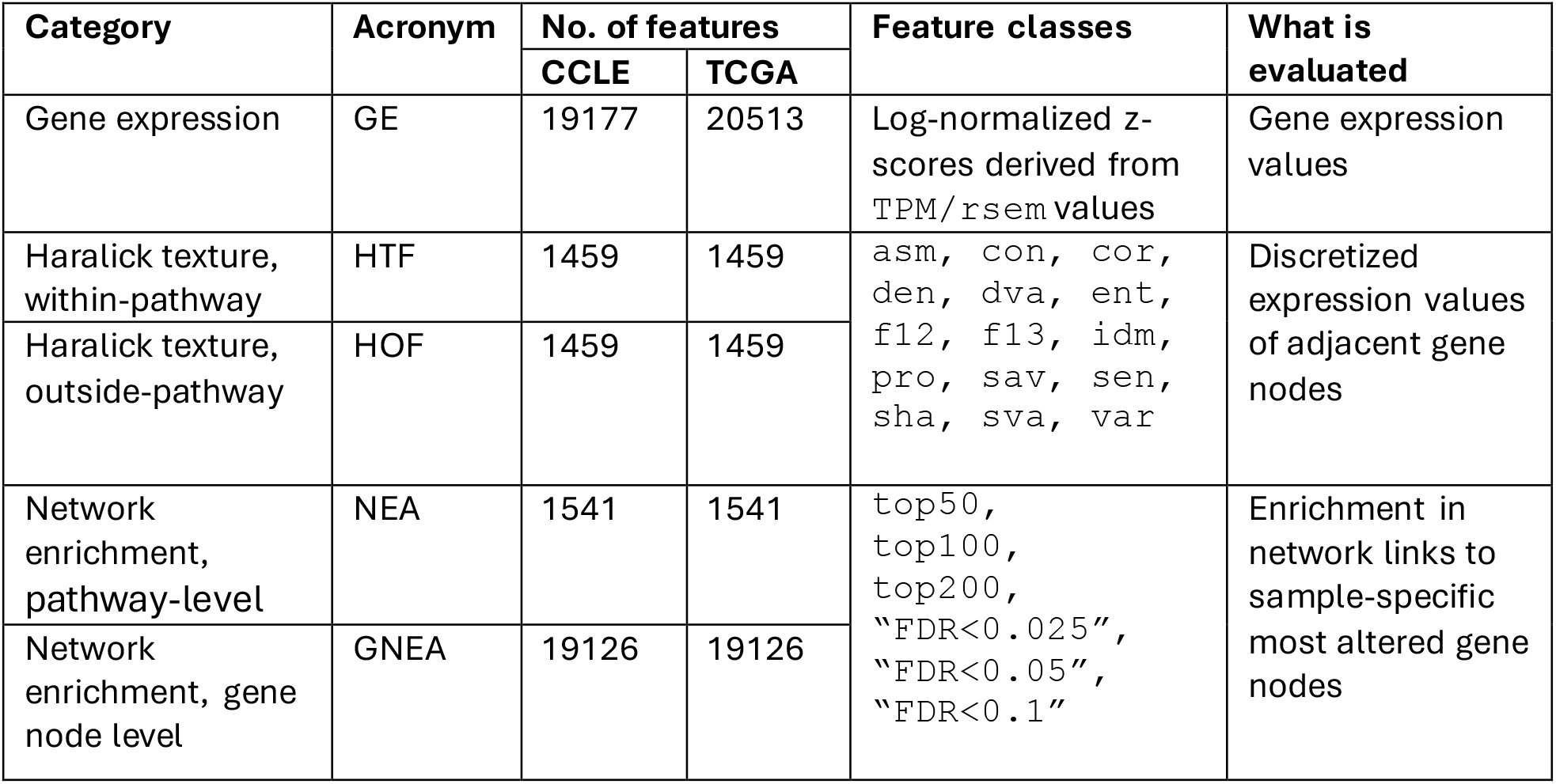
Features used in comparative analyses.

| Category | Acronym | No. of features |  | Feature classes | What is evaluated |
| --- | --- | --- | --- | --- | --- |
|  |  | CCLE | TCGA |  |  |
| Gene expression | GE | 19177 | 20513 | Log-normalized z-scores derived from TPM/rsem values | Gene expression values |
| Haralick texture, within-pathway | HTF | 1459 | 1459 | asm, con, cor, den, dva, ent, f12, f13, idm, pro, sav, sen, sha, sva, var | Discretized expression values of adjacent gene nodes |
| Haralick texture, outside-pathway | HOF | 1459 | 1459 |  |  |
| Network enrichment, pathway-level | NEA | 1541 | 1541 | top50, top100, top200, "FDR<0.025", "FDR<0.05", "FDR<0.1" | Enrichment in network links to sample-specific most altered gene nodes |
| Network enrichment, gene node level | GNEA | 19126 | 19126 |  |  |

First, the t parameter, which controls the number of discretization bins 2^*t*^, was optimized by minimizing FDR levels across the three CCLE drug-screen analyses. Among the practical values *t*={2,3,4,5}, *t*=3, i.e. discretization into 2^3^=8 equally sized bins, was used in all subsequent analyses.

For each alternative gene- and pathway-level profile, correlations with drug-sensitivity profiles were calculated for three large-scale anti-cancer *in vitro* screens. The CCLE collection was split into cohorts by tumor site of origin, of which the three most abundant, breast, lung, and lymphoid, were used as predictors in univariate linear models of drug response. Comparative performance, while controlling the false discovery rate at three significance levels, is presented in Fig. 2 (see also Supplementary File 1) as fractions of significant correlates in each feature class. When comparing feature classes, the analysis should not be interpreted only through the central tendency of the boxplots. In a downstream multivariate setting, only a small subset of the strongest individual features would usually be selected. Therefore, high-performing outliers are potentially relevant, if they are not isolated artifacts and if their occurrence is consistent with the broader distribution of the corresponding feature class. Overall, the best HTF/HOF features (`con`, `sav`, and `var`) frequently reached the performance levels of the best NEA/GNEA features – especially for lung-derived cell lines, while the bulk of their distributions was also broadly comparable. Also, certain classes of HTF and HOF features demonstrated high similarity between, suggesting redundancy and comparable statistical performance (Fig. 1F). After considering the dendrogram and the performance in the sensitivity tests described above, we chose the features `con`, `den`, `f13`, `sav`, `sha`, and `var` from both the HTF and HOF classes for further detailed investigation.

**Figure 2.**
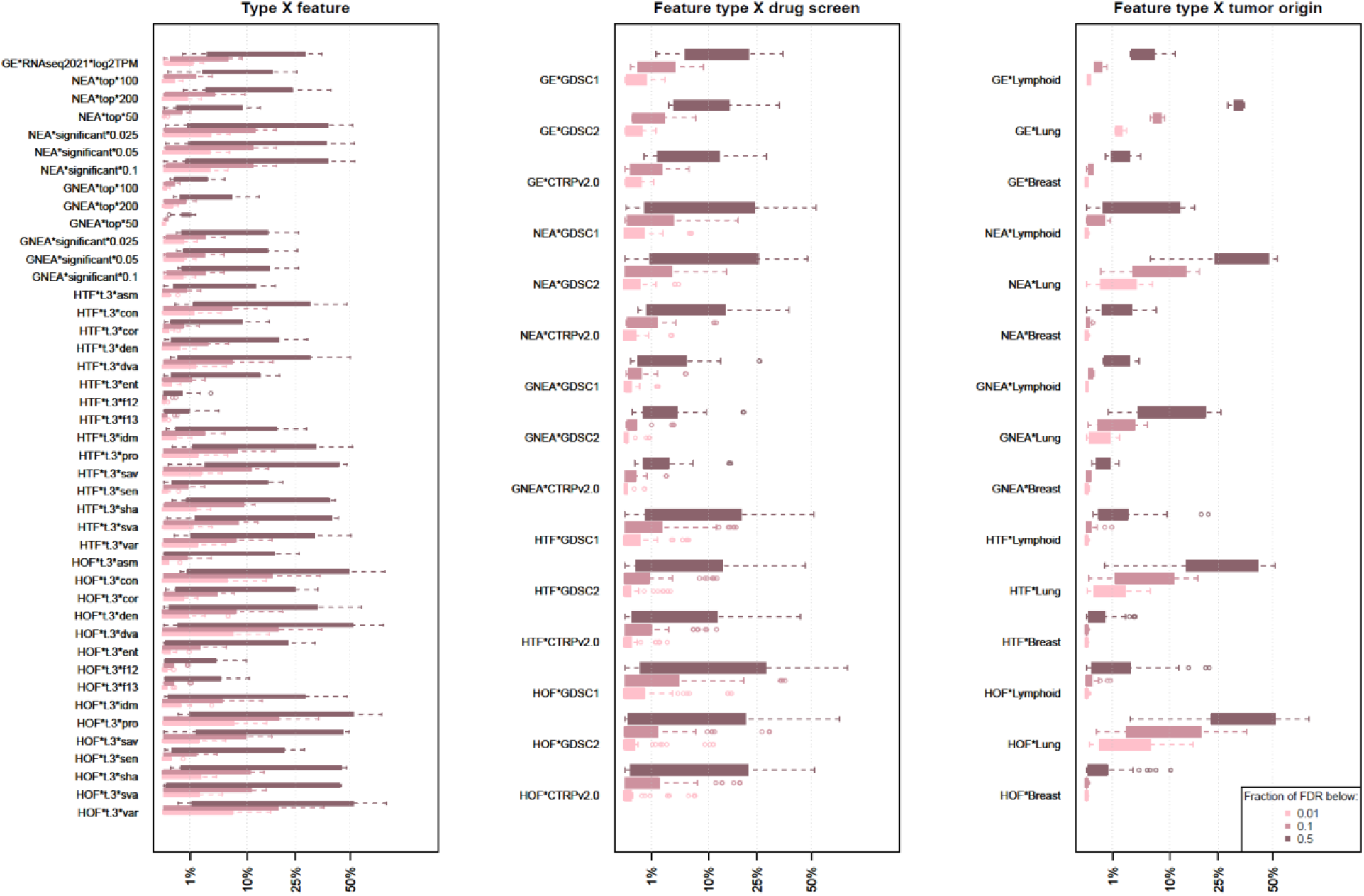
Performance of feature classes on CCLE data collection. The plots represent levels of performance achieved in univariate models of drug sensitivity from *in vitro* screens. The boxes combine model p-values for all drugs available in the three screens, adjusted for multiple testing for each drug separately. Feature performance was evaluated by the ability to explain drug response and summarized as fractions of models in each box with false discovery rates below the cutoffs FDR < {0.01, 0.1, 0.5}. The whiskers extend to the most extreme data points that are no more than 1.5 times the box length away from the box. More distant points are shown as outliers. The box hinges correspond to the first and third quartiles. The notches extend to +/-1.58 IQR/sqrt(n), which roughly corresponds to a 95% confidence interval for the difference between two medians (Mcgill et al. 1978).

### Preservation of drug sensitivity correlates between *in vitro* drug screens

The analysis above tested the overall ability of feature classes to explain drug response, with false positive rates addressed only through formal multiple-testing correction. For practical use, however, correlates should remain stable under changes in screening methodology, batch structure, and biological composition. Robustness was therefore evaluated using three independent drug-screen datasets: GDSC1, GDSC2, and CTRP v.2. In total, 138 drugs were shared by at least two screens. For each shared drug, agreement between two screens was estimated as the Spearman rank correlation between drug-feature association p-values. Because tumor site of origin strongly affects molecular phenotypes and drug response, site was included as a covariate in a two-factor model, with the feature effect estimated as a separate term. In parallel, univariate models were applied to seven site-specific cell-line subsets.

In contrast to the initial analysis, the cross-screen comparison showed that pathway-based features were more reproducible than original GE features (Fig. 3): the GE correlates agreed substantially less well between screens. Although pathway-based feature classes varied at increasingly stringent rank thresholds, differences between them were usually not significant. Most importantly, the algorithmically simpler HTF/HOF features performed not worse than NEA/GNEA.

**Figure 3.**
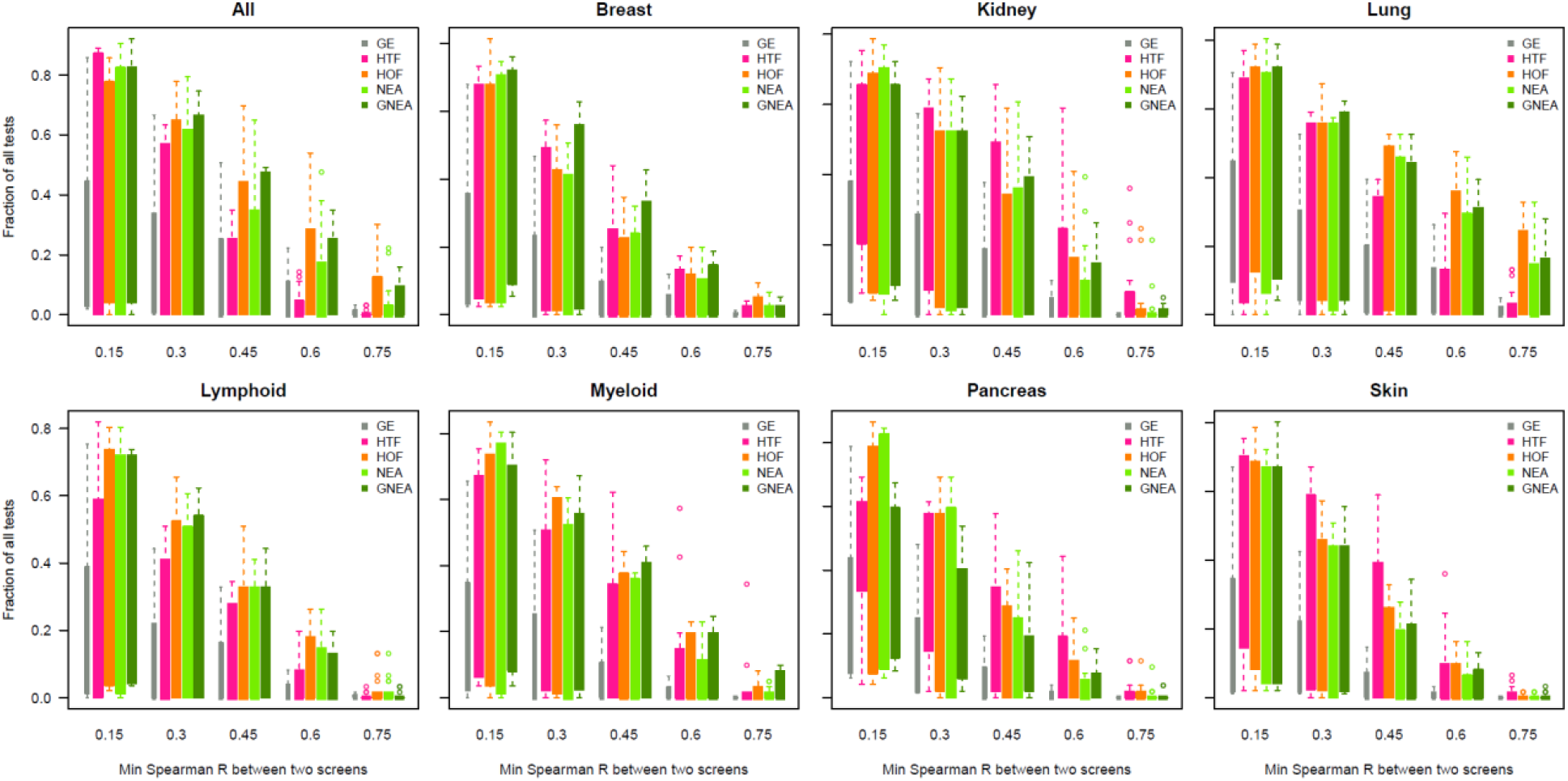
Consistency of drug-feature correlates between *in vitro* drug screens. For each drug tested in two screens, Spearman rank correlation coefficients were calculated between p-values of drug-feature correlation in the two screens. The bins combine tests of all drugs with all feature classes of each category (see Table 1). The axes display fractions of drug-feature correlates preserved between the two screens at levels no worse than *r* > *c*, where *c* is one of the five raising correlation levels *R* ∈ {0.15…0.75}. The compared p-values were derived from covariate models accounting for the site origin (label “All”) and from site-specific univariate models (“Breast”, “Kidney” etc.).

### Preservation of correlates between *in vitro* and clinical patient profiles

Analysis of TCGA clinical cohorts required a more complex statistical design. Because disease stage at diagnosis strongly influences patient survival, “stage” was included as a covariate in Cox proportional-hazards models together with “drug” and “feature” terms. Statistical significance of the main “feature” term indicated an association with survival irrespective of treatment. By this criterion, certain HTF and HOF features, such as `con` and `var`, outperformed other feature classes (Fig. 4A).

**Figure 4.**
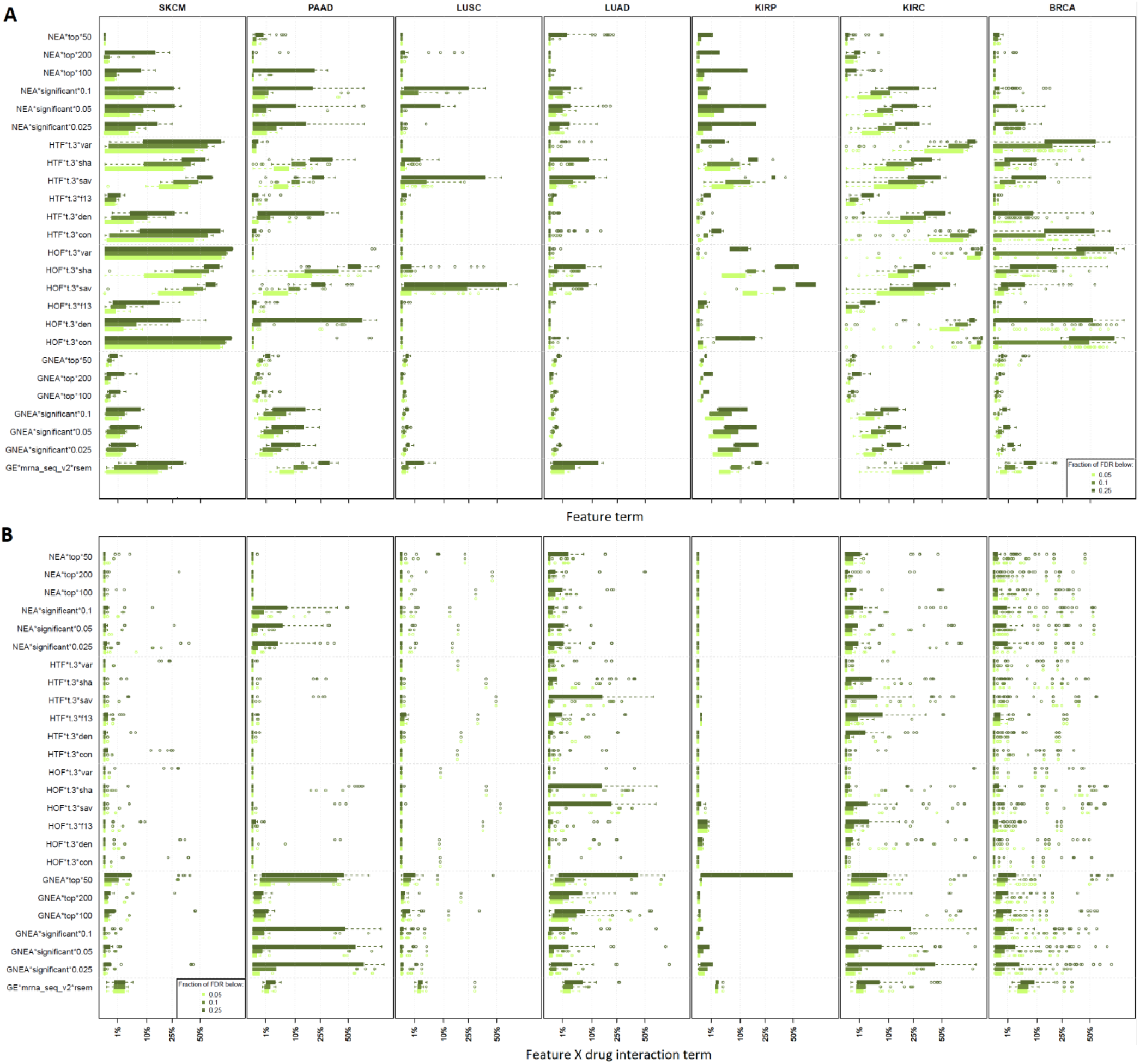
Performance of feature classes on TCGA cohort data. The plots represent levels of performance achieved in Cox proportional-hazards models reporting the effects of features, drug treatment, their interactions, and clinical covariates. The boxes combine tests across all drugs and all follow-up intervals available in the cohorts. A. Fractions of all models in which the adjusted p-value for the main term “feature” was below an FDR threshold {0.05, 0.1, 0.25} (Benjamini and Hochberg 1995). B. Same as A, for the “drug × feature” interaction term.

The practical objective, however, is to identify features that are potentially predictive of drug response. Such a property should appear as a survival difference among patients who received a given drug, but not among patients who did not receive it. To evaluate this effect, a “drug × feature” interaction term was included in the model. A significant interaction indicates that higher or lower feature levels are associated with altered survival under treatment. The model is illustrated by the four-stratum Kaplan-Meier plots in Figure 5: survival differed by feature level among treated patients but not among untreated patients. With respect to the interaction term, HTF/HOF features generally performed below NEA/GNEA, although selected features were competitive in the LUAD and KIRC cohorts (Fig. 4B).

**Figure 5.**
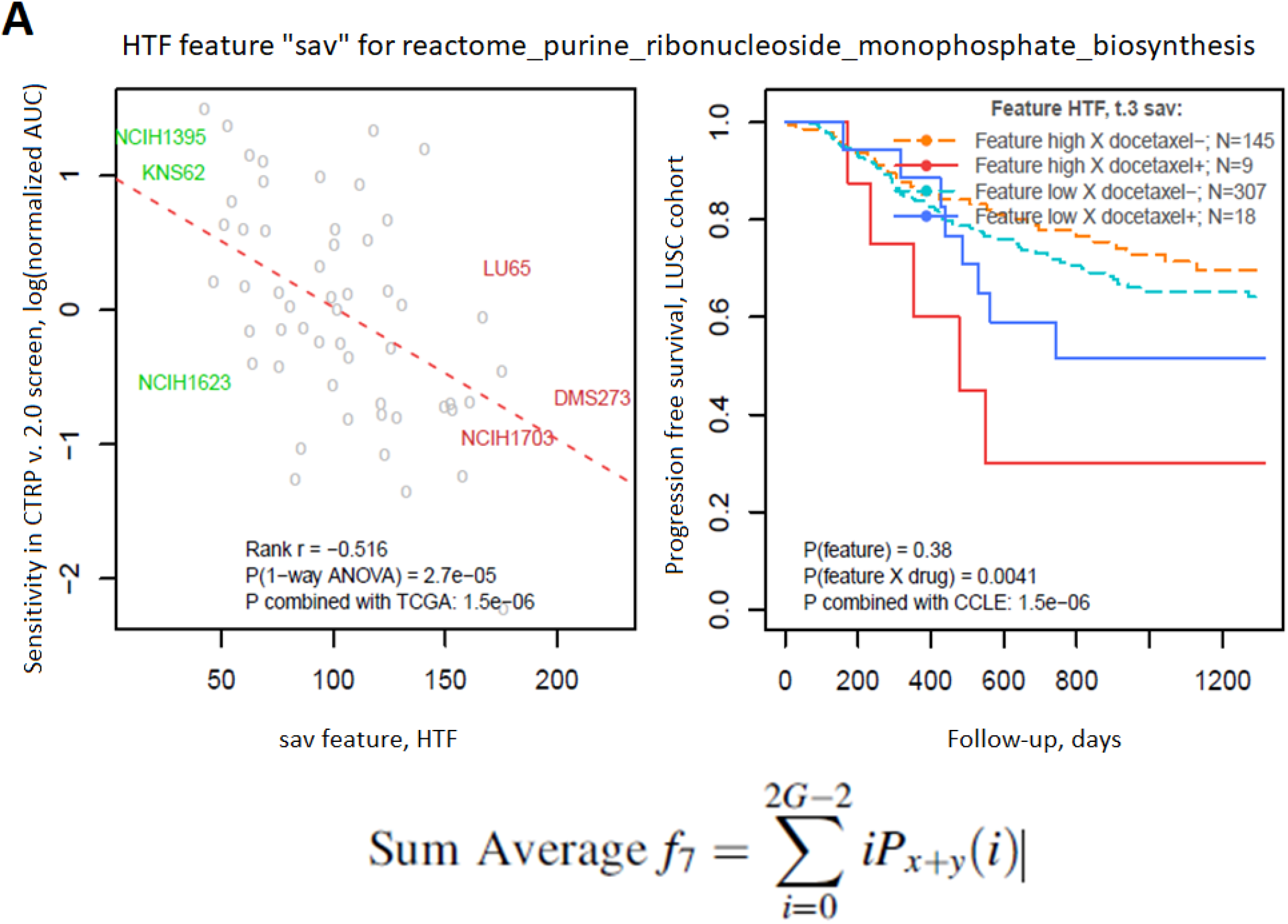
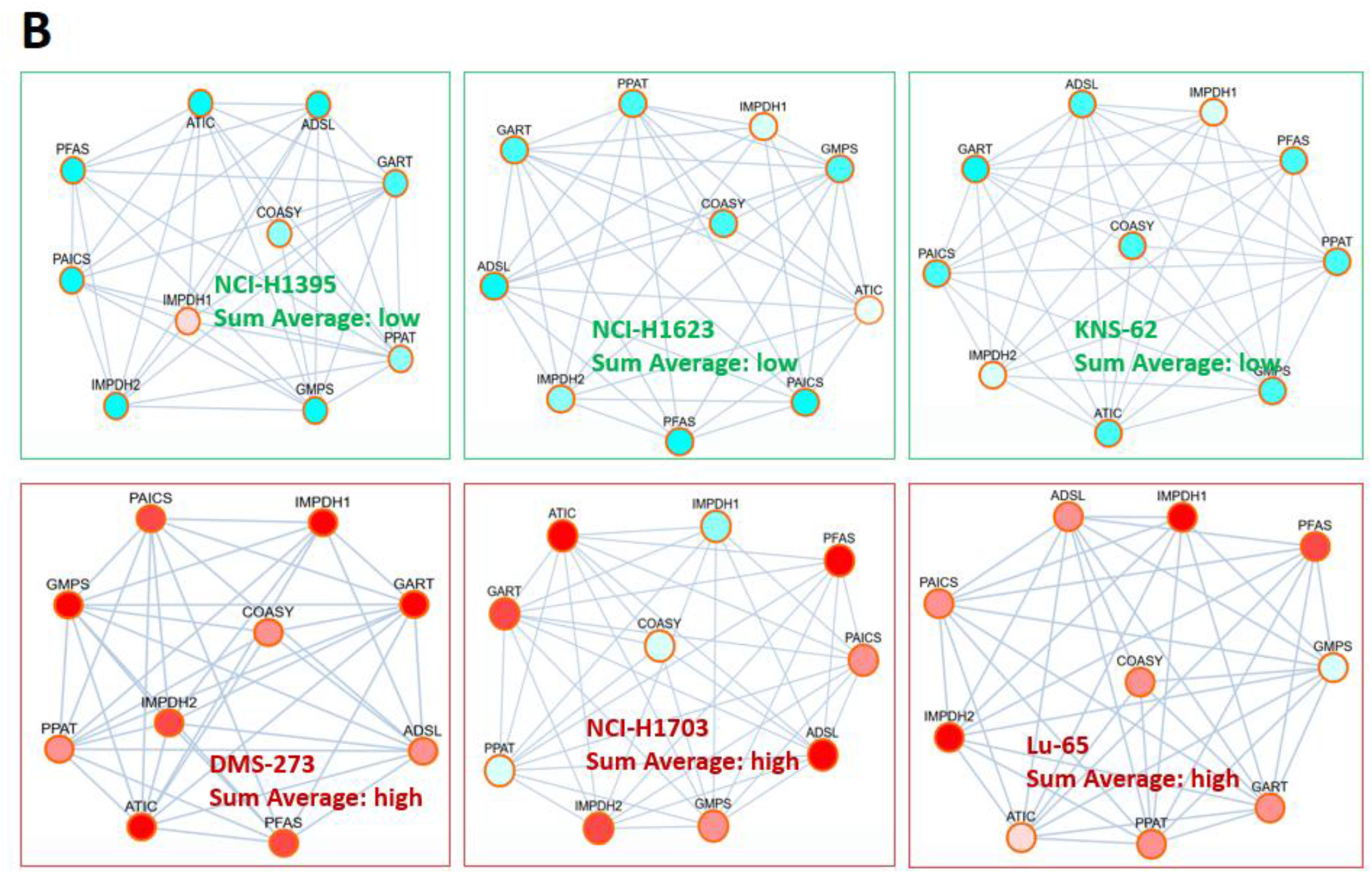
Preservation of drug-to-feature interactions between *in vitro* and clinical datasets. A. Regression of sensitivity to docetaxel in the CTRP v. 2.0 screen on the `sav` feature for the purine synthesis pathway (left), and the corresponding analysis of patient survival depending on administration of the same drug and the `sav` value for the same pathway (right). Details for the six highlighted cell-line samples are shown in B. B. Example “activity-connectivity” patterns where higher `sav` values were achieved in cell lines where activity was synchronously high in pairs of adjoint nodes.

Using this framework, we then addressed the most relevant and challenging question: to what extent are drug-feature correlates detected in CCLE preserved in TCGA survival analyses? Features of the same methods and classes were generated from gene expression by applying the same pipeline to CCLE and TCGA datasets. The analysis was restricted to drugs represented in both *in vitro* screens and clinical treatment records. After requiring that a drug had been administered to at least seven patients in a cohort, up to ten drugs were available per cohort and 20 drugs were available in total. The same drug could occur in up to five cohorts, but each cohort was analyzed separately. The p-value distributions indicated that the best HTF and HOF features performed at approximately the same level as NEA features (Fig. 6; Supplementary Files 2 and 3).

**Figure 6.**
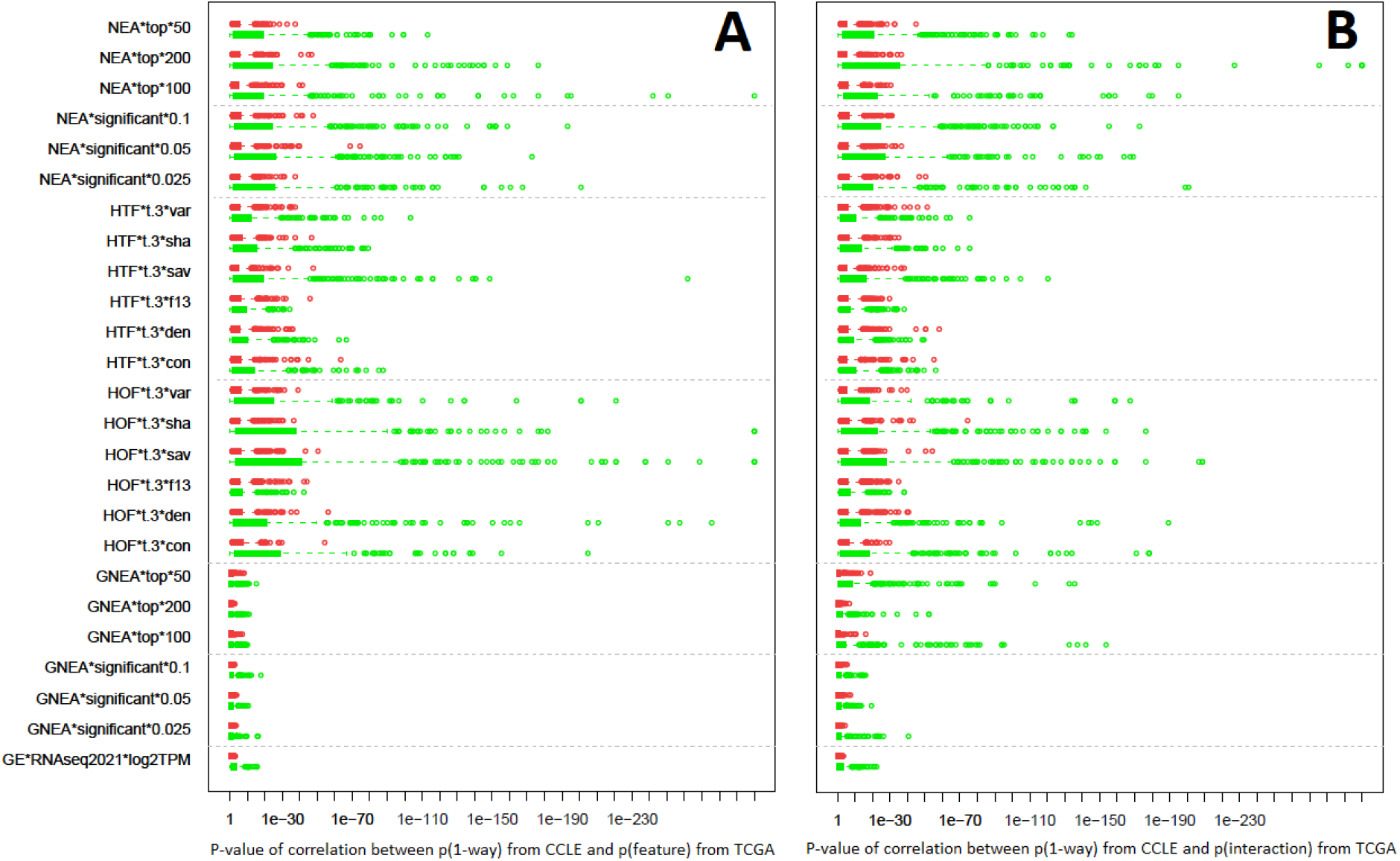
Agreement between drug sensitivity *in vitro* and in the clinic. The plots represent levels of agreement between the CCLE *in vitro* screen analysis and p-values from Cox proportional-hazards models of overall survival in the six TCGA cohorts. The boxes combine tests across all drugs and all follow-up intervals available in the cohorts. To account for effect directions for each drug, model p-values were multiplied by regression coefficient signs. Agreement between signed p-values from CCLE and TCGA analyses was then quantified using Spearman rank correlations (green). To control false positive rates, p-value distributions from random permutations of the same data profiles are shown in red.

If the same drug-feature relationship is biologically relevant *in vitro* and in patients, the direction of association should also be preserved. This was tested by matching effect sizes from the *in vitro* p.1x model to interaction terms from the survival model. Across all feature categories (GE, NEA, GNEA, HTF, and HOF), association signs in the actual data were preserved 2.2-to 6-fold more often than in randomly permuted sample and patient vectors (Bonferroni adjusted p-values from Fisher’s exact test ranged from 0.028 to 0.3×10^-24^).

## Methods

### Network

The global network *N*(*V*,*E*) consists of gene nodes represented as vertices *V* and functional links represented as edges *E*. The network used in this study was created from STRING (version 12; (Szklarczyk et al. 2017) by selecting the top 999,589 edges ranked by STRING confidence score, and contained 19,126 HUGO gene-symbol nodes.

### Pathway-specific matrices

For any given pathway *p*, we extract from network *N*(*V*,*E*) a sub-graph *P*(*V*_*p*_,*E*_*p*_):*V*_*p*_ ∈*V, E*_*p*_ ∈*E*. No self-loops or duplicate edges were allowed, all edges were undirected, and edge weights were ignored after the top-edge selection step.

The global network N is very dense and set *E*_*p*_ usually contains many more edges between genes of *p* than the respective formal versions provided by e.g. KEGG or Reactome.

Under the default HTF option, *E*_*p*_ includes only edges connecting member genes of *p* with each other. The extended HOF option also considers the so-called edge boundary of the pathway subgraph, that is, outgoing edges in which one of the two connected genes does not belong to the pathway:

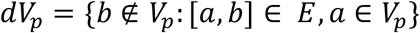

The gene expression matrix for |*V*_*p*_ | genes in *S* samples 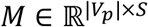 is retrieved from a transcriptomics dataset. The values of *M* are discretized into 2^*t*^ = *G: t* ∈{2,3,4…} intensity levels to become sample-specific attributes of gene nodes *g*. This results in a matrix 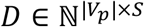 of discrete values *e*_*gs*_ ∈{0…*G-*1}: *g* ∈*V*_*p*_, *s* ∈ *S*. From this matrix, a so-called gray-level co-occurrence matrix *GLCM* = ℕ^×*G*^ is generated; its elements *GLCM*_*i*,*j*_ summarize co-occurrence of intensities *i* and *j* in adjoint nodes *A*_*a*,*b*_ = 1, ∀*E*_*a*,*b*_ ∈ |*E*_*p*_|:*e*_*a*_=*i*, *e*_*b*_=*j*. After division by the sum of its elements, *GLCM* becomes a probability matrix, from which the texture features are calculated at the next step.

### Haralick texture features

The R package `wvtool` (Kobayashi et al. 2015; https://github.com/pywood21/wvtool) implemented calculation of the original set of textural features proposed by Haralick et al. 1973, plus two features (`sha`, `pro`) suggested by Albregtsen (1995). The feature acronyms to the right of each formula are used throughout the present paper:

Angular Second Moment: 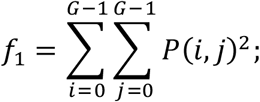 (**asm**)

Contrast: 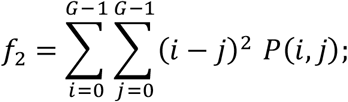 (**con**)

Inverse Difference Moment: 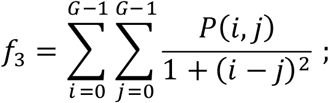(**idm**)

Enthropy: 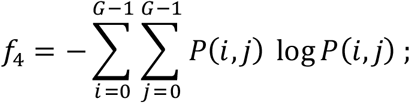 (**ent**)

Correlation: 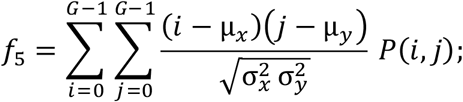 (**cor**)

Variance: 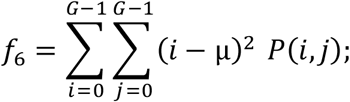 (**var**)

Sum Average: 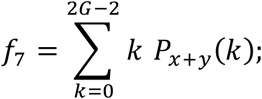 (**sav**)

Sum Entropy: 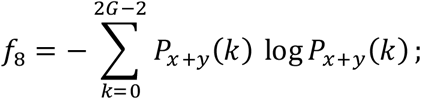 (**sen**)

Difference Entropy: 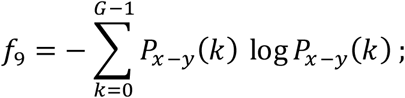 (**den**)

Difference Variance: 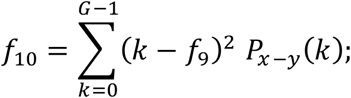 (**dva**)

Sum Variance: 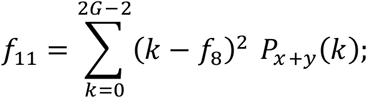 (**sva**)

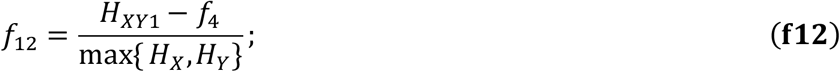

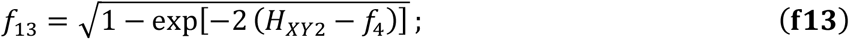

Shade: 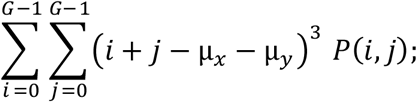 (**sha**)

Prominence: 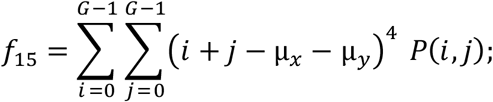 (**pro**) where *y* and *x* correspond to rows and columns of GLCM and intensity levels *i* and *j*, so that *P*(*i, j*) is the (*i, j*)th element of a GLCM, *G* being the number of gray levels: {*i*,*j*}∈1…G;

The marginal-probability values are obtained by summing up the rows and columns of GLCM:

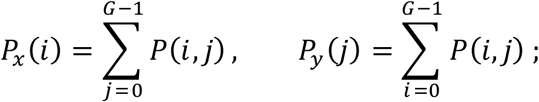

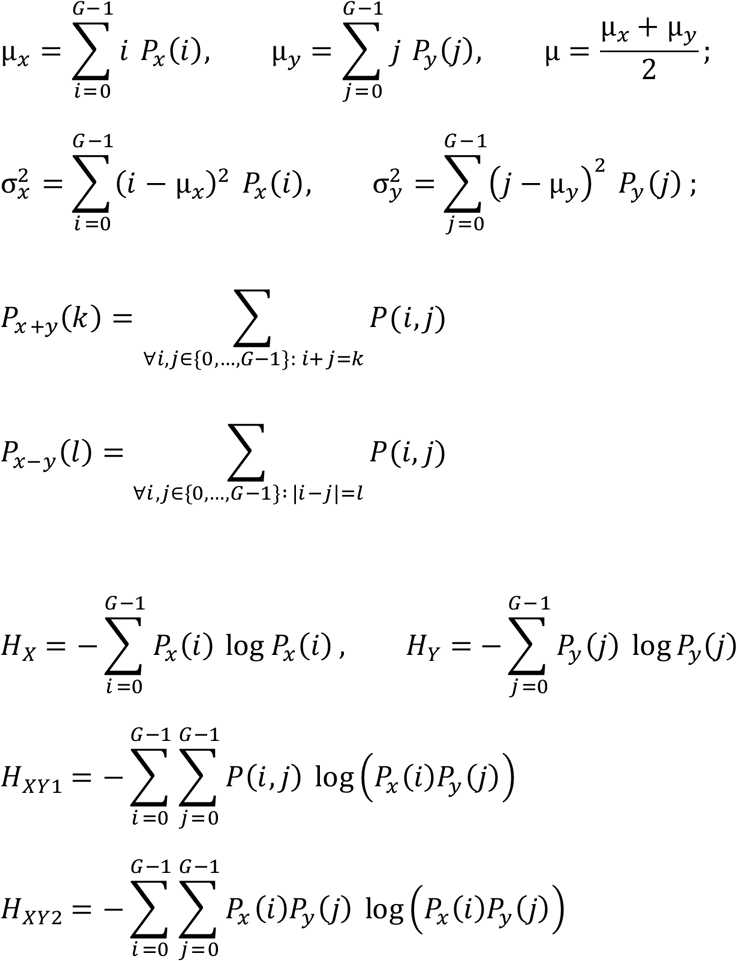

### Modifications of original code

We inspected and optimized code of the `wvtool` package according to the formulas above; these changes did not alter the numerical output. The main modification was the calculation of GLCMs from graphs with arbitrary node degrees. Before GLCM construction, the expression matrix can optionally be normalized by row, normalized by column, or left unnormalized.

The functions `glcm.net` and `haralick.net` for calculating HTF and HOF features, together with the wrapper function `haralick.matrix`, are included in R package `NEArender v. 2.0` alongside the NEA functions described below. Execution in parallelized mode is enabled.

### Network enrichment analysis

The NEA algorithm (Jeggari and Alexeyenko 2017) enables enrichment analysis in terms of network links connecting two gene sets of interest: an altered gene set (AGS), most often characterizing an experiment or a biological condition, and a functional gene set (FGS), such as a pathway or a Gene Ontology term. The binomial statistic 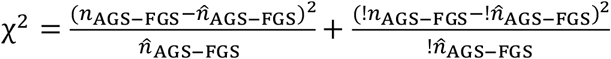 evaluates relative abundance of network links between any genes of AGS and any genes of FGS; links within AGS and within FGS are ignored. The respective numbers expected by chance are calculated as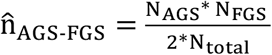, where *N*_*AGS*_ and *N*_*FGS*_ sum the node degrees of member genes, and *N*_*total*_ is the number of edges in the whole network. The second term of χ^2^ statistic represents the rest of the global network, where “!” denotes “other than”. The non-negative statistic values are converted to the normally distributed NEA z-score and signed, so that *z*<0 indicates depletion.

### Transcriptomics data

The CCLE and TCGA transcriptomics datasets described below were used in parallel as original expression features and for generating expression-based HTF, HOF, NEA, and GNEA features.

#### Cell line dataset

The Cancer Cell Line Encyclopedia (CCLE) collection (Ghandi et al. 2019) contains distinct and systematically typed cell lines, of which 1,377 were profiled by RNA-seq (dataset file CCLE_expression_full.csv downloaded in 2021 from https://depmap.org). TPM expression values were log-transformed and normalized as *z*-scores. Cell lines known to originate from breast (N=62), kidney (N=44), lung (N=184), pancreas (N=65), skin (N=84), lymphoid (N=152), and myeloid (N=61) tumors were used as site-specific subsets in the univariate statistical model. The first five of these sites were matched to TCGA cohorts as described below. In parallel, 1,246 cell lines from these and other well-represented sites (N>75), namely bone, head and neck, bowel, esophagus/stomach, and CNS/brain, were included in the analysis using a multi-site statistical model with site of origin as a covariate.

Drug sensitivity in the three alternative *in vitro* screens, CTRP v. 2.0 (Seashore-Ludlow et al. 2015), GDSC1 and GDSC2 (Yang et al. 2013), was evaluated from curves over 6 or 8 sequential dilution points and expressed as the area under the curve (AUC). For the analysis in this study, AUC values were transformed toward normality by inverse normal transform (Beasley et al. 2009) across the tested cell lines, for each drug separately.

#### TCGA data

The Cancer Genome Atlas datasets were downloaded from https://www.cbioportal.org/datasets for the following cohorts:

##### Breast

BRCA (Koboldt et al. 2012)

##### Lung

adenocarcinoma (LUAD, Collisson et al. 2014) and squamous cell cancer (LUSC, Hammerman et al. 2012).

##### Kidney

clear renal cell carcinoma (KIRC, Creighton et al. 2013) and papillary renal-cell carcinoma (KIRP, Cancer Genome Atlas Research Network 2016).

##### Pancreas

ductal adenocarcinoma (PAAD, Cancer Genome Atlas Research Network and Cancer Genome Atlas Research Network 2017)

##### Skin

cutaneous melanoma (SKCM, Cancer Genome Atlas Research Network 2015).

In the analysis of conserved correlates, these five TCGA categories were matched to the corresponding CCLE sites listed above.

The RNA-seq files “`rsem_zscores_ref_all_samples`” provided by the cBioPortal team were used as the corresponding transcriptomics datasets.

Follow-up profiles for relapse-free or progression-free survival and overall survival were available and were evaluated in parallel. Because the relevant timeframe can depend on cancer aggressiveness and treatment course, alternative follow-up intervals were defined as one-quarter, one-half, and the full available follow-up period.

### Drug effect models

#### Linear models of cell line response

The univariate statistical model for CCLE profiles estimated significance of feature effects *β*_*f*_ on cell line sensitivity of each specific site of origin of drug *d*:

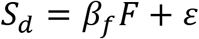

In the multi-site statistical model, the site of origin was included in the model as a covariate *C*:

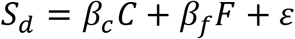

After accounting for the effect of site, the significance of the feature effect was estimated and reported with the respective p-value for *β*_*f*_.

### Patient survival models

Because treatment response and molecular features can depend strongly on tumor stage and other clinical variables, drug-feature associations must be estimated with covariate adjustment. To avoid unrealistically complex models, only prognosis-associated covariates were included: `ajcc_pathologic_tumor_stage` for each TCGA cohort and, additionally for BRCA, `er_status_by_ihc`, `pr_status_by_ihc`, and `her2_status_by_ihc`. A Cox proportional-hazards model was fitted for each feature-drug combination:

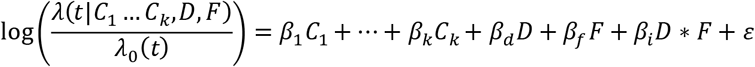

Together with covariates *C*, the model included the main effects of drug *D* and feature *F* as well as the interaction term *D×F*. The drug variable was encoded as any exposure, regardless of timing and combination therapies. Drug names were harmonized across patient records. A significant main drug effect can be interpreted as an overall treatment-associated survival difference irrespective of the feature. Conversely, a significant feature effect indicates that the feature is associated with survival regardless of treatment. The interaction term *D×F* is therefore central to testing drug-feature associations, because it asks whether the relationship between feature level and survival differs between treated and untreated patients. Main effects of feature and drug were allowed but were not required. A significant interaction should indicate that survival differs by feature level specifically among patients who received the drug, whereas untreated patients show no comparable feature-dependent survival difference.

All survival analysis calculations were done using R package `survival`. The statistical significance of the model terms was estimated with `coxph` using continuous feature vectors. For visualization the Kaplan-Meier curves the feature levels were binarized at a cutoff.

#### Adjustment for multiple testing

The false discovery rate (FDR) values reported in Figs. 2 and 4 were estimated with the Benjamini-Hochberg method (Benjamini and Hochberg 1995) for each tested drug separately.

## Discussion

Compared with raw gene-expression features, the proposed Haralick-type texture features and higher-order features produced pathway-level profiles with markedly reduced dimensionality and better preservation of drug-response correlates across datasets. Their performance was broadly comparable to NEA/GNEA rather than clearly superior to it. The main advantage of the new approach is therefore not an absolute gain in robustness, but its technical simplicity: it avoids explicit construction of altered gene sets, operates directly on quantitative node values, and can be extended by defining additional graph-level texture features. This makes the method a practical complement to NEA/GNEA and a flexible source of candidate variables for downstream predictive modelling.

The observational nature of TCGA treatment data is a major limitation of this study. Although tumor stage was included in the clinical analysis, administration of specific drugs could still be confounded by unmeasured clinical conditions and unknown factors. Neither the proportional-hazards assumption nor e.g. the requirements for Mendelian randomization analysis were fully met, making unbiased estimation of drug-by-feature effects unfeasible. Nonetheless, the analytical setup served the comparative purpose of the study: evaluating whether different methods and method-specific feature classes preserve *in vitro* correlates in clinical data. The absence of randomized clinical controls was also addressed by random permutation tests. By this criterion, preservation of the correlates discovered *in vitro* was better for pathway-level NEA and HTF features than for gene-level GNEA and, especially, original gene-expression profiles (Fig. 6).

A practical advantage of Haralick features over NEA is that the magnitude, and therefore potential sensitivity, of several Haralick features did not depend on pathway size or on the total/mean pathway node degrees. This was particularly clear for the best-performing features `sha`, `sav`, and `var`, and also for `con`, but not for `den` or `f13` (Supplementary File 4). In contrast, absolute NEA values correlated with pathway size and total node degree, while, as expected from NEA normalization, no correlation with mean node degree was observed.

Haralick and NEA features share one important property: both account for sample-specific gene activity. However, NEA considers only most altered genes and ignores activity of pathway genes unless they are also included in the AGS. When pathway-level features were calculated with NEA, normalization by node degree was required because node degree sums vary across sample-specific AGSs. Haralick features do not use AGSs, so node-degree variability across samples is absent for a fixed pathway and normalization not required for within-pathway comparisons.

Normalization would, however, be needed for direct comparisons between pathways or between whole networks with different degree distributions. Analyses such as those of Barker-Clarke and co-authors, which compared whole networks with different topologies and degree distributions, would therefore benefit from explicit degree adjustment.

However, even node-degree normalization in NEA may not fully account for higher-order topological properties. The relative independence of HTF/HOF features from such properties is therefore valuable, although cross-pathway comparisons of raw HTF/HOF magnitudes are not necessarily interpretable. Further development of the method should address this need with a suitable normalization technique.

The ability to vary or extend the set of textural features is another advantage of the Haralick approach and remains an avenue for future mechanistically oriented network analysis work.

## Abbreviations

AGS: altered gene set
AUC: area under the curve
BRCA: breast invasive carcinoma
CCLE: Cancer Cell Line Encyclopedia
CNS: central nervous system
CTRP: Cancer Therapeutics Response Portal
ER: estrogen receptor
FDR: false discovery rate
FGS: functional gene set
GDSC: Genomics of Drug Sensitivity in Cancer
GE: gene expression
GLCM: gray-level co-occurrence matrix
GNEA: gene-level network enrichment analysis
GO: Gene Ontology
HER2: human epidermal growth factor receptor 2
HOF: Haralick outside-pathway feature
HTF: Haralick texture feature
IQR: interquartile range
KEGG: Kyoto Encyclopedia of Genes and Genomes
KIRC: kidney renal clear cell carcinoma
KIRP: kidney renal papillary cell carcinoma
LUAD: lung adenocarcinoma
LUSC: lung squamous cell carcinoma
NEA: network enrichment analysis
ORA: overrepresentation analysis
PAAD: pancreatic adenocarcinoma
PCA: princoipmalponent analysis
PH: proportional hazards
PR: progesterone receptor
RNA-seq: RNA sequencing
SKCM: skin cutaneous melanoma
ssGSEA: single-sample gene set enrichment analysis
TCGA: The Cancer Genome Atlas
TPM: transcripts per million

