## Supplementary figures and images for "Textural features for pathway-level representation of omics data in biological networks"

### Supplementary File 1

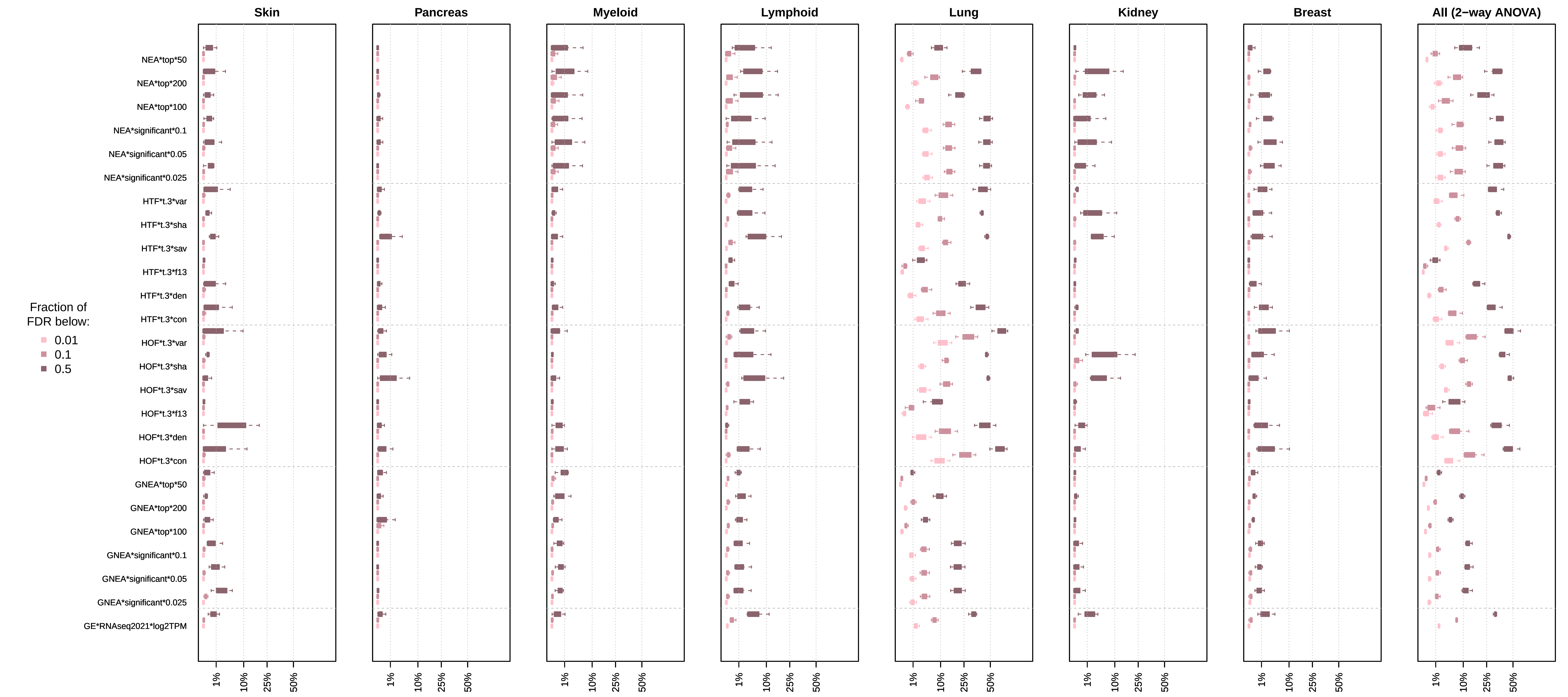

### Supplementary File 2

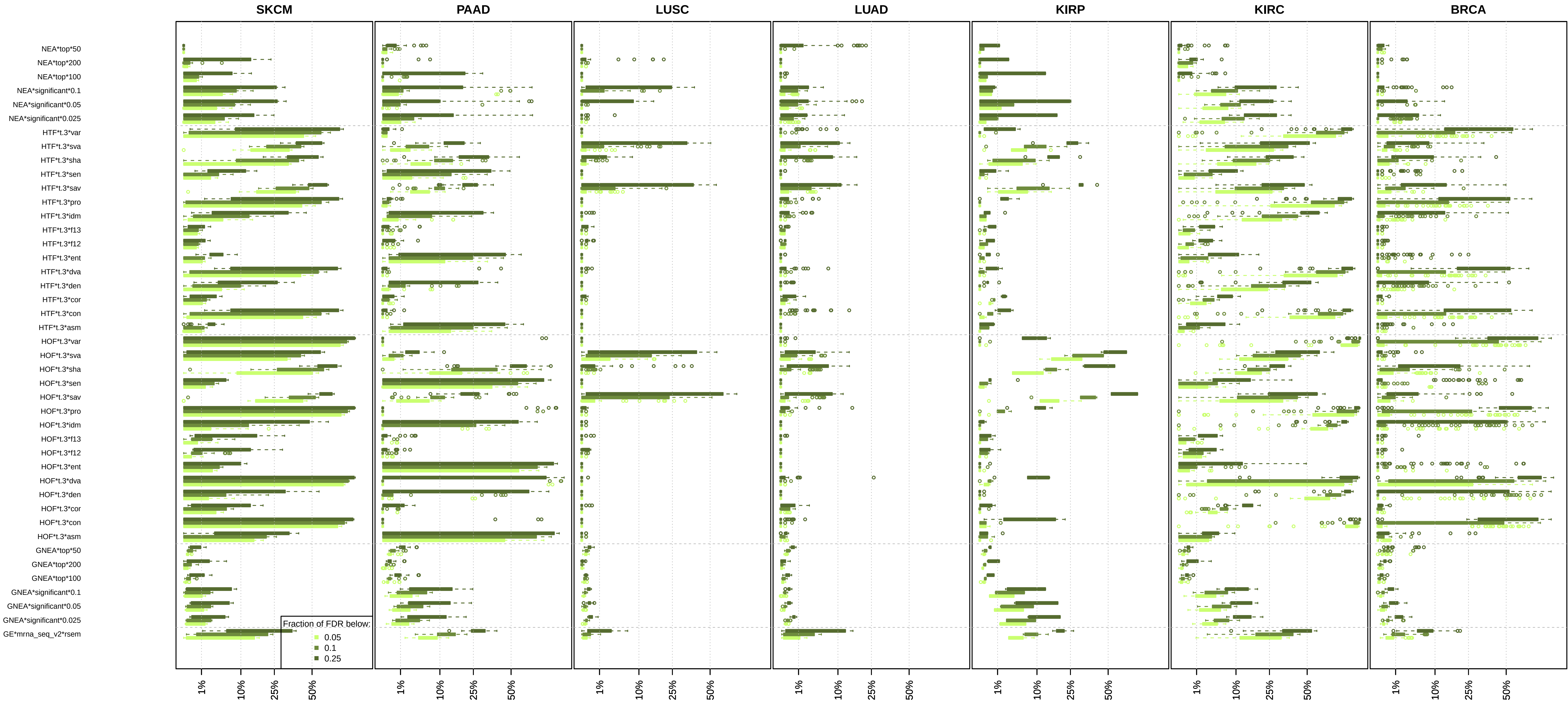

### Supplementary File 3

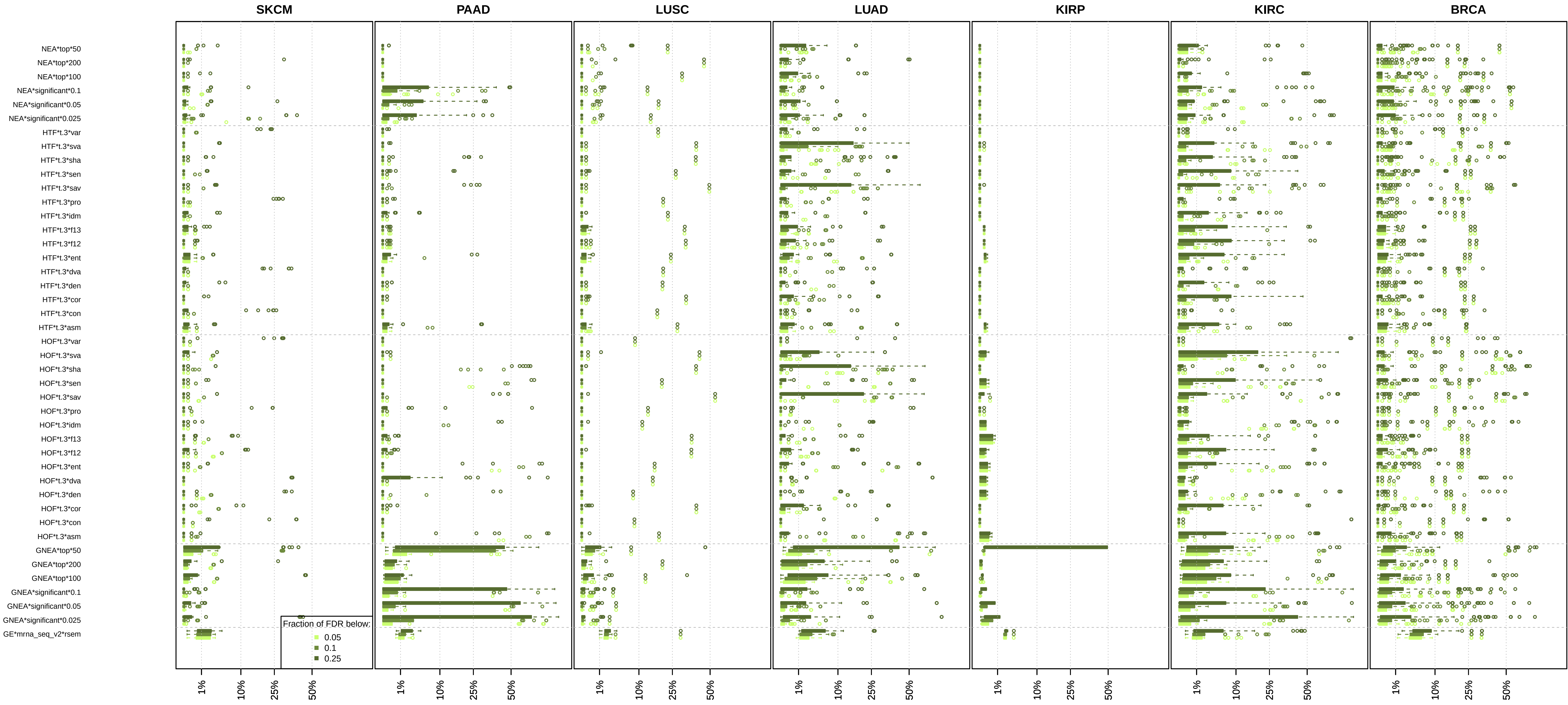

### Supplementary File 4

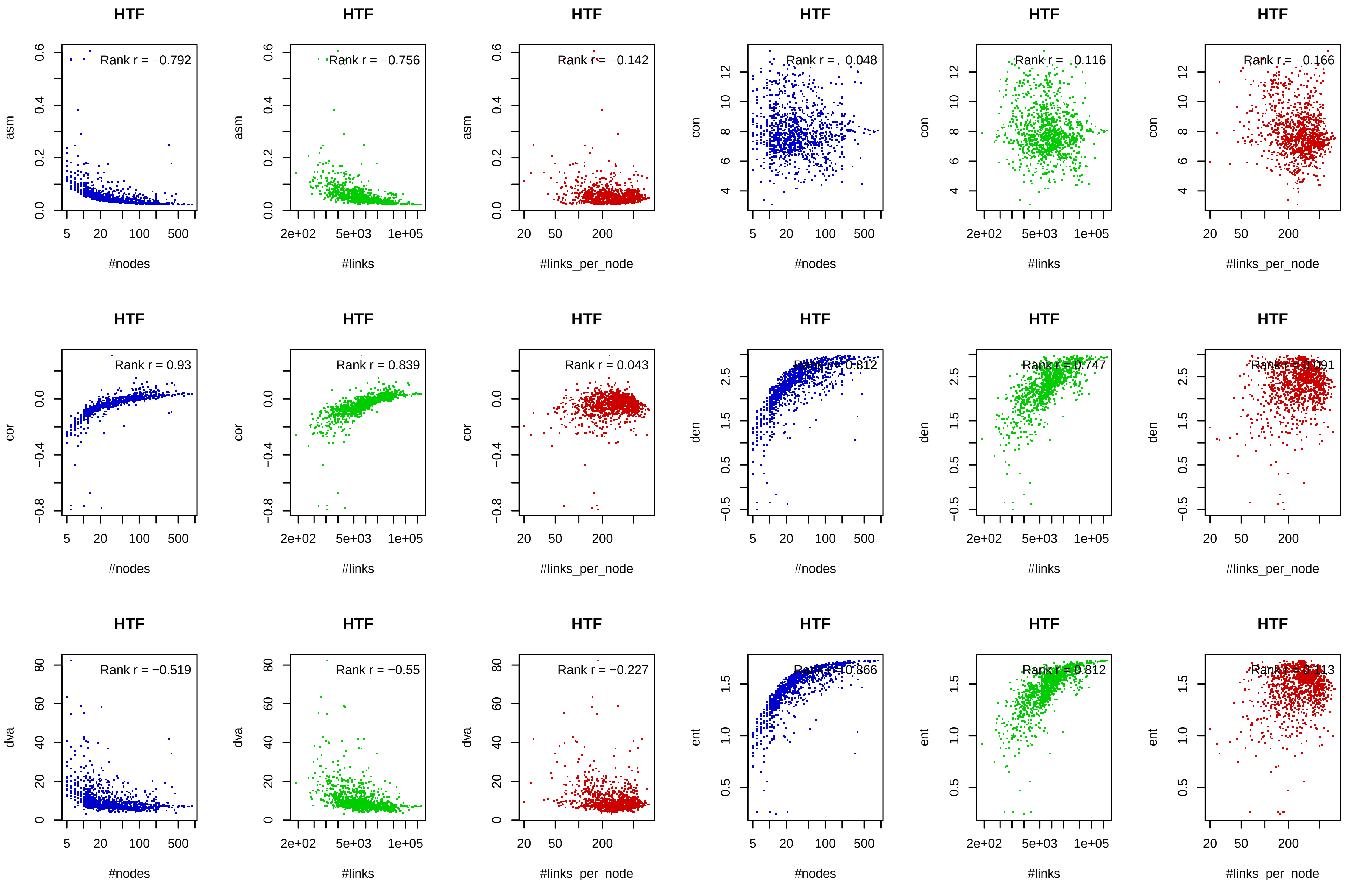

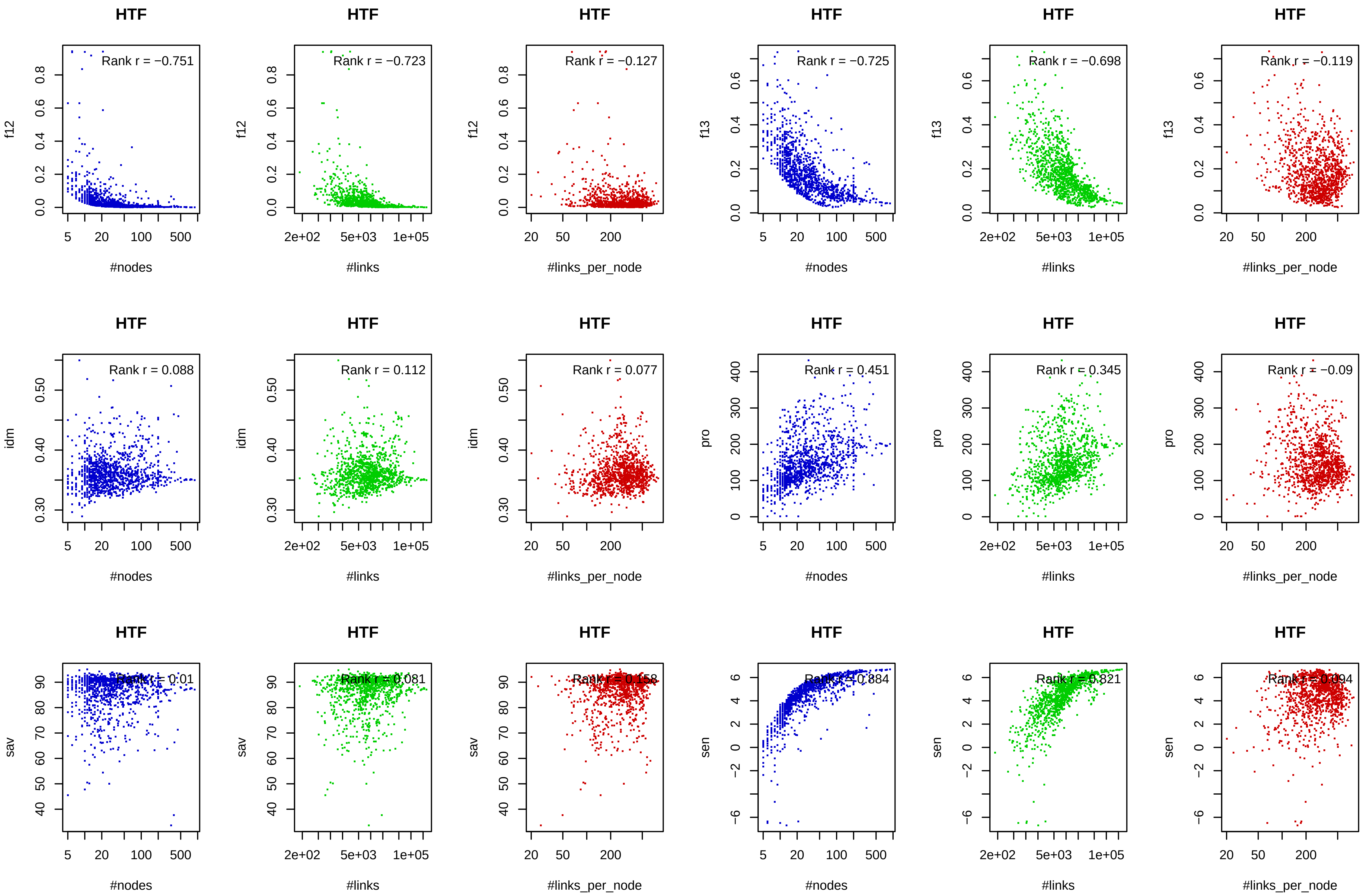

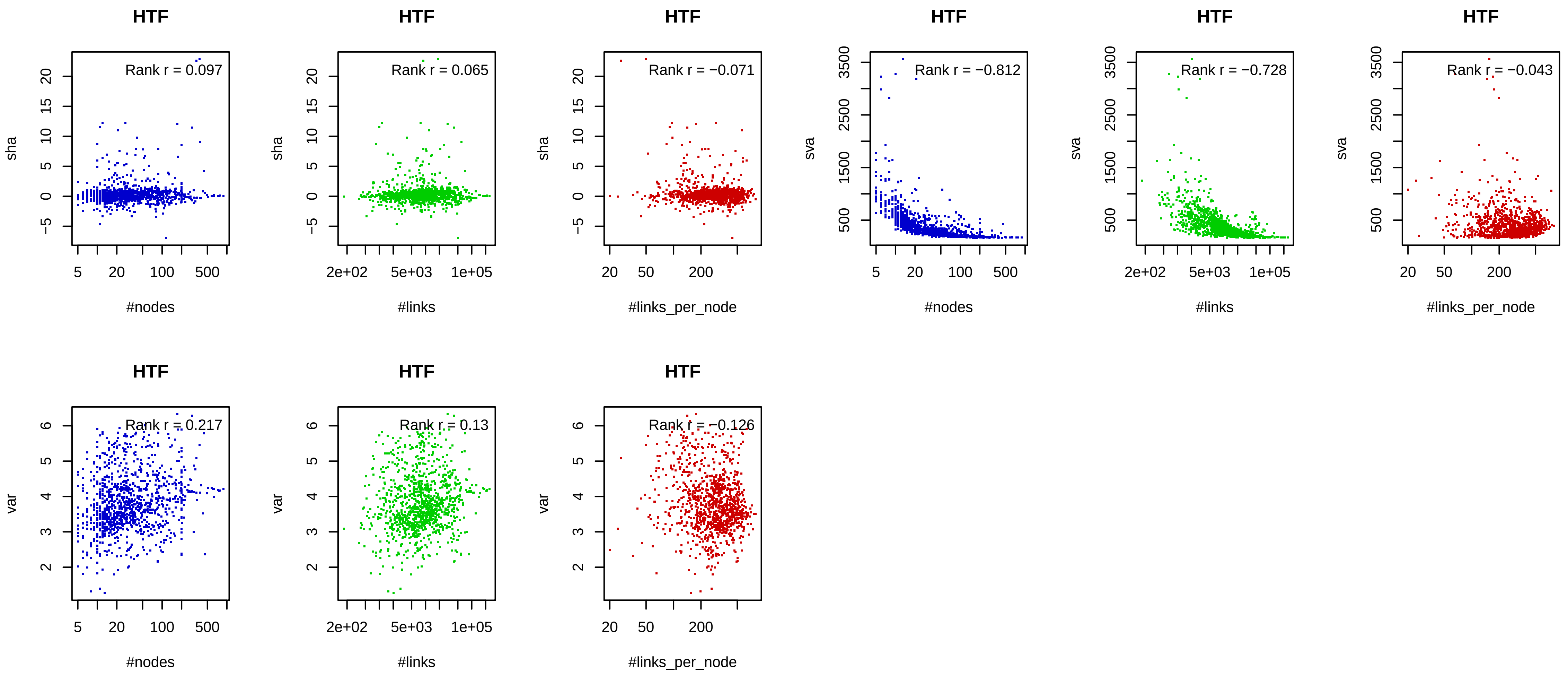

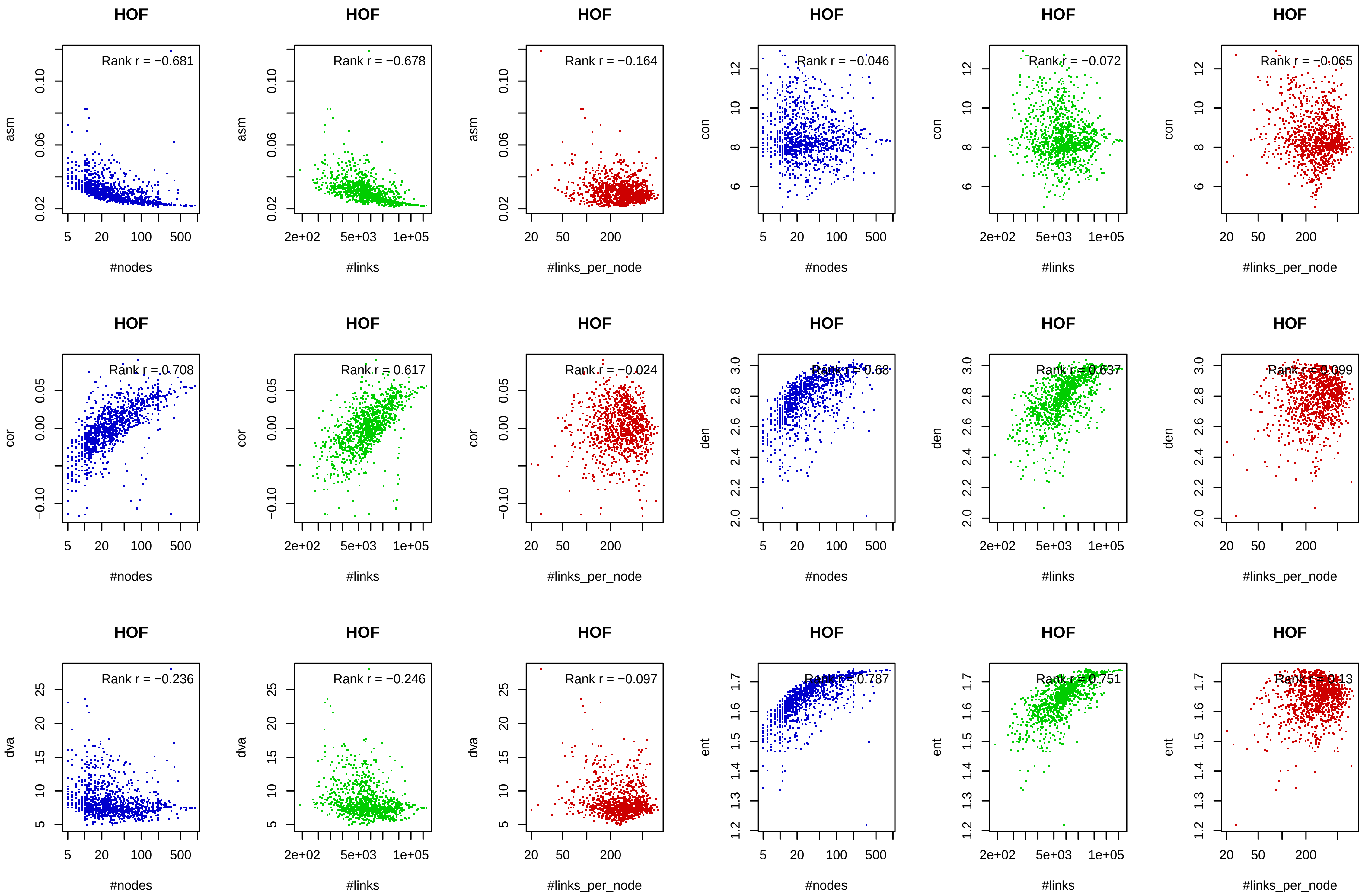

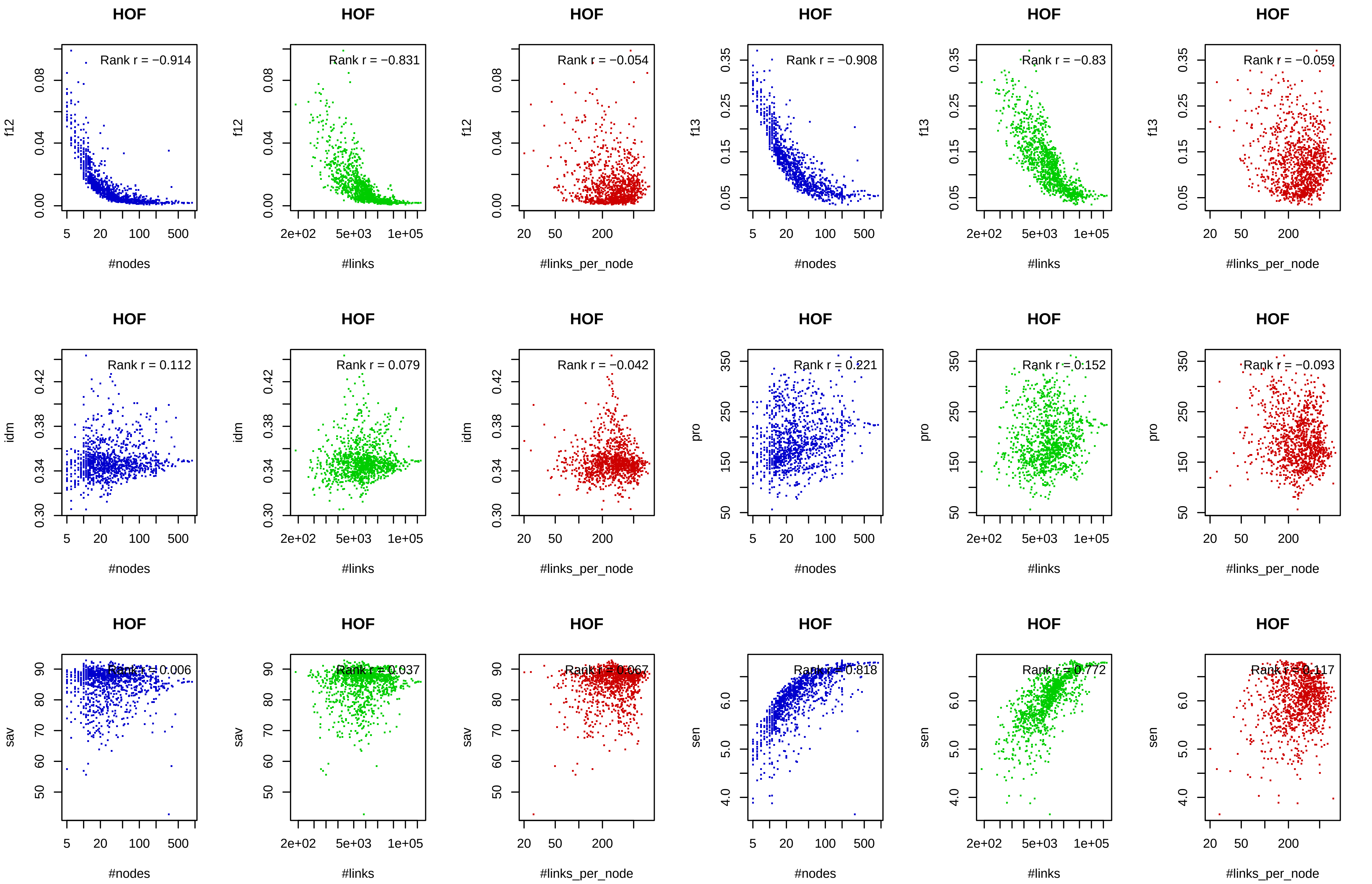

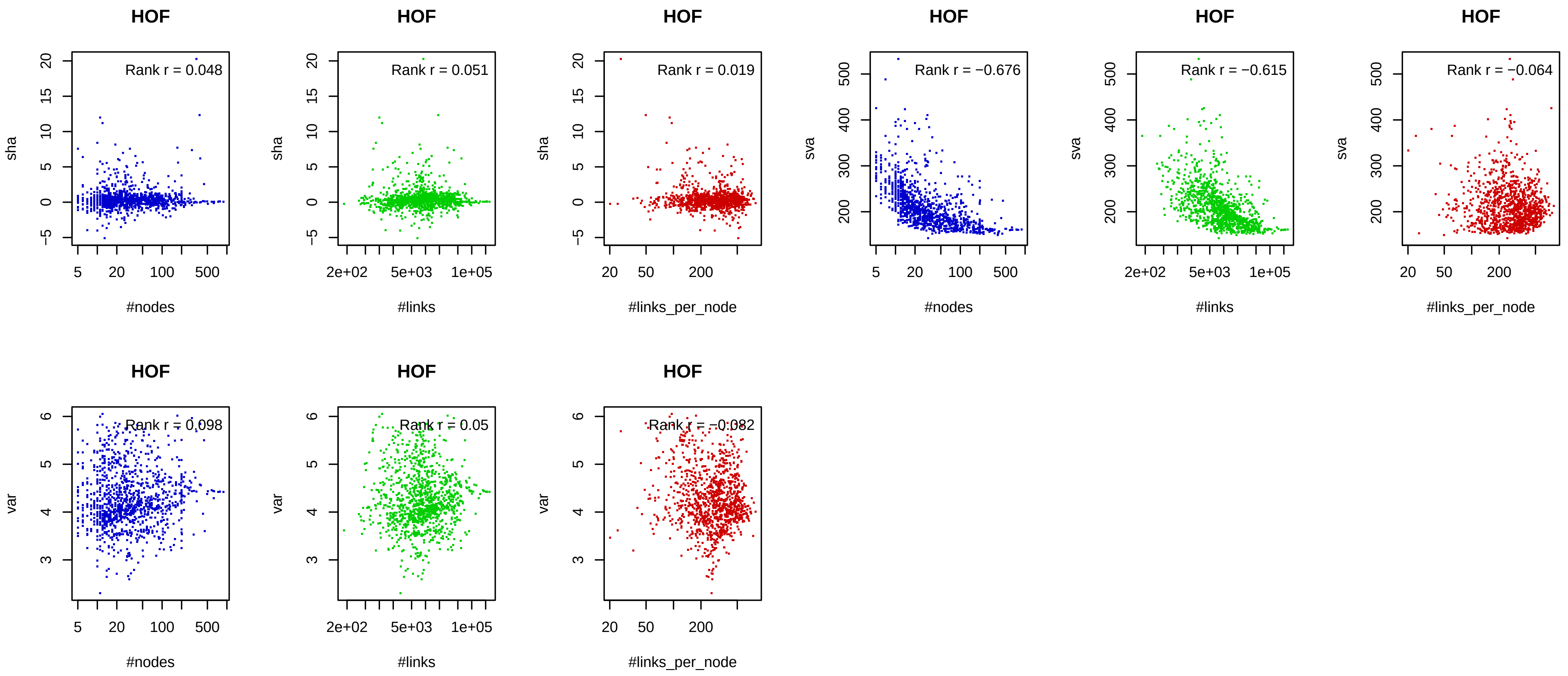

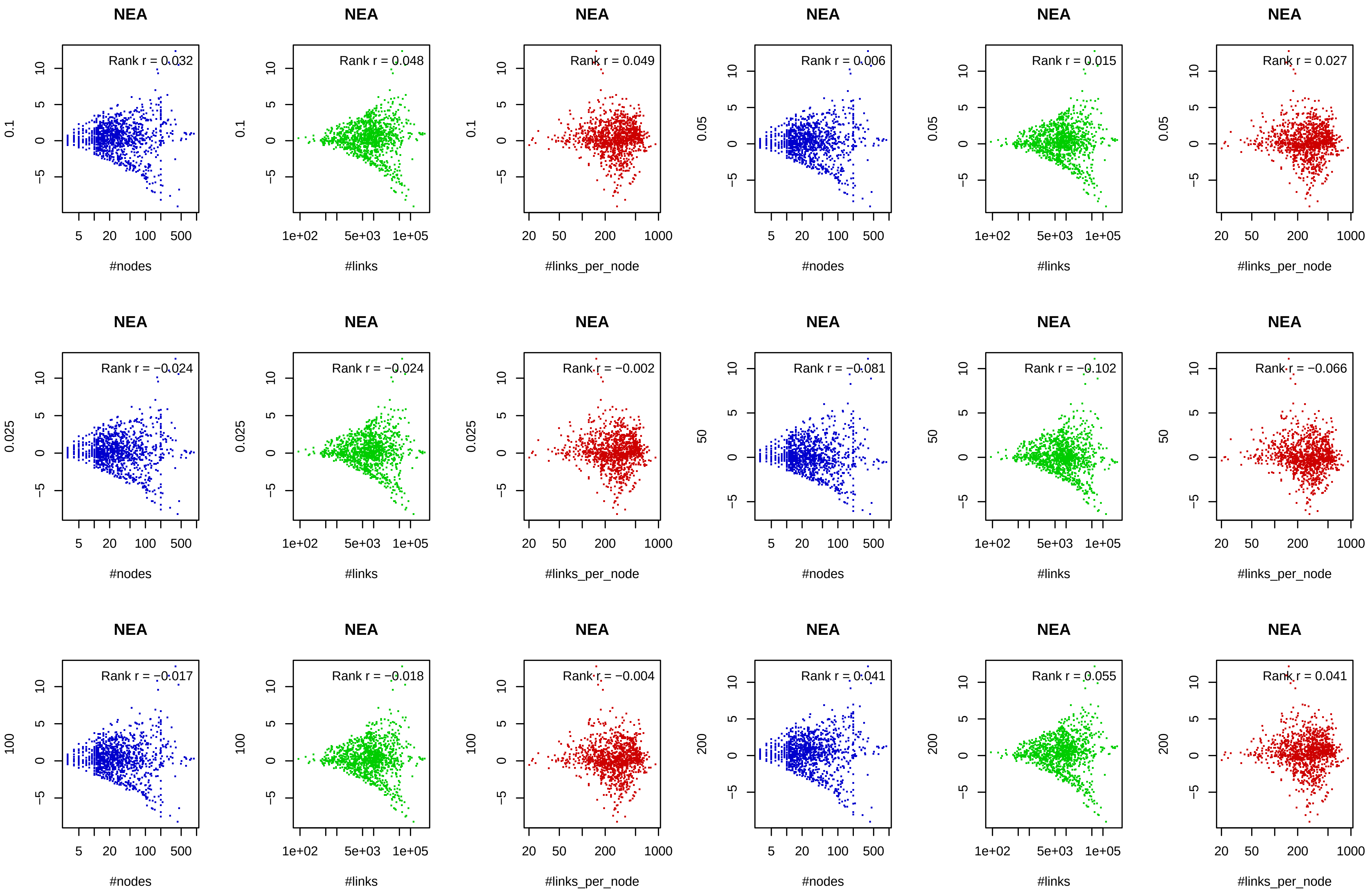
